# The anterior Hox gene *ceh-13* and *elt-1/GATA* activate the posterior Hox genes *nob-1* and *php-3* to specify posterior lineages in the *C. elegans* embryo

**DOI:** 10.1101/2021.02.09.430385

**Authors:** John Isaac Murray, Elicia Preston, Jeremy P. Crawford, Jonathan D. Rumley, Prativa Amom, Breana D. Anderson, Priya Sivaramakrishnan, Shaili D. Patel, Barrington Alexander Bennett, Teddy D. Lavon, Felicia Peng, Amanda L. Zacharias

## Abstract

Hox transcription factors play a conserved role in specifying positional identity during animal development, with posterior Hox genes typically repressing the expression of more anterior Hox genes. Here, we dissect the regulation of the posterior Hox genes *nob-1* and *php-3* in the nematode *C. elegans*. We show that *nob-1* and *php-3* are co-expressed in gastrulation-stage embryos in cells that previously expressed the anterior Hox gene *ceh-13*. This expression is controlled by several partially redundant transcriptional enhancers. These enhancers require *ceh-13* for expression, providing a striking example of an anterior Hox gene positively regulating a posterior Hox gene. Several other regulators also act positively through *nob-1/php-3* enhancers, including *elt-1/GATA*, *ceh-20/ceh-40/Pbx*, *unc-62/Meis*, *pop-1/TCF*, *ceh-36/Otx* and *unc-30/Pitx*. We identified defects in both cell position and cell division patterns in *ceh-13* and *nob-1;php-3* mutants, suggesting that these factors regulate lineage identity in addition to positional identity. Together, our results highlight the complexity and flexibility of Hox gene regulation and function and the ability of developmental transcription factors to regulate different targets in different stages of development.

**Author Summary:** Hox genes are critical for head-to-tail patterning during embryonic development in all animals. Here we examine the factors that are necessary to turn on two posterior Hox genes*, nob-1* and *php-3*, in the nematode worm*, C. elegans*. We identified six new transcription factors and three enhancer regions of DNA that can activate expression of *nob-1/php-3*. Unexpectedly, these activating transcription factors included *ceh-13*, an anterior Hox gene, and *elt-1*, a regulator of skin development that is briefly expressed in many cells that do not adopt skin fates, including the cells that express *nob-1*. Furthermore, the cellular defects we observed in *ceh-13* and *nob- 1;php-3* mutant embryos indicate that the early embryonic functions of these Hox genes help determine the identity of cells as well as their position within the embryo. Our findings identify new roles for Hox genes in *C. elegans* and emphasize the ability of transcription factors to contribute to the diversification of cell types and the adoption of specific cell types at different phases of embryonic development.

## Introduction

Hox genes are conserved transcription factors famously expressed in specific positions along the anterior-posterior axis during animal development that specify axial position. Mutations in Hox genes cause a wide variety of developmental defects in both model organisms and humans. Hox gene regulation is complex and includes both transcriptional and post-transcriptional control. In most animals, Hox genes are found in genomic clusters, and their expression along the A-P axis is collinear with their genomic position, and often show “posterior dominance,” where posterior Hox genes repress the expression of more anterior Hox genes (Reviewed in (1–3)).

The genome of the nematode *Caenorhabditis elegans* encodes a single set of six Hox genes on Chromosome III, loosely organized into three degenerate “clusters” that each contain two adjacent genes (4–6,3). The cluster containing *lin-39* and *ceh-13* is inverted relative to Hox clusters in other organisms as *ceh-13*, a homolog of *HoxA1/labial,* is downstream of *lin-39,* a homolog of *HoxA4/Deformed*(7–9). In larval stages, these genes are expressed in specific positions along the A-P axis and regulate both positional differences in cell fate and function, similar to their homologs in other animals (10–18), and directly regulate terminal fates (19).

Three of these, the anterior Hox gene homolog *ceh-13/HOX1* and the posterior Hox gene homologs *nob-1/HOX9-13* and *php-3/HOX9-13,* are also expressed in early embryogenesis, during gastrulation (5,20–22). *ceh-13* mutants have severe defects in anterior (head) morphology(21), while mutants lacking both *nob-1* and *php-3* have severe posterior (tail) defects(5). Partial cell lineage tracing of *ceh-13* and *nob-1*;*php-3* mutants identified defects in cell position but not in division patterns, leading to the hypothesis that these genes regulate positional identity, rather than lineage identity(5,6,21).

Intriguingly, despite their apparently opposite roles in anterior vs posterior morphogenesis, both *ceh-13* and *nob-1* are expressed in overlapping posterior lineages during gastrulation and both require the Wnt pathway for this expression(23, 24). *ceh-13* is transiently expressed in the progeny of 7 of the 8 posterior sister cells derived from the (largely ectoderm- producing) AB blastomere at the 24-cell stage(20). *nob-1* is expressed 1-2 cell cycles later, in the posterior daughters or grand-daughters of four of these *ceh-13*-expressing cells(22). In addition, both *ceh-13* and *nob-1* are expressed in the posterior daughter of the intestinal blastomere E (“Ep”). This raises the question of whether *ceh-13* regulates *nob-1* in these lineages. Expression of both factors at later stages is regulated by feedback mechanisms; early *ceh-13* activity is required for later ceh-13 expression(24), and early *nob-1* negatively regulates later *nob-1* expression through the microRNA *mir-57*(22).

Here we analyzed the *cis-*regulatory control of *ceh-13* and *nob-1* expression in the early embryo. We find that nob-1 expression is regulated by several partially redundant distal enhancers, including at least three that drive overlapping patterns during gastrulation. Two of these enhancers are positively regulated by *ceh-13*, while a third appears to be regulated by a *ceh-13*-independent mechanism. We further identified the GATA family transcription factor *elt-1*, previously known as a specifier of epidermal fate, as one of several additional positive regulators of *nob-1* expression. Detailed analysis of cell positions and cell division timing identifies both position defects and defects in cell division patterns and timing in *ceh-13* and *nob-1*/*php-3* mutants. This positive regulation of a posterior Hox gene by an anterior Hox gene suggests a novel role of Hox genes in early lineage specification.

## Results

### The anterior Hox gene *ceh-13* and the posterior Hox gene *nob-1* are expressed sequentially in gastrulating embryos

To better understand the expression dynamics of the *ceh-13, nob-1,* and *php-3* Hox genes during embryogenesis (Figure 1), we collected 3D time-lapse movies (∼1.5 minute temporal resolution) of embryos expressing either *ceh-13, nob-1* or *php-3* tagged with GFP at their C-termini at the endogenous locus by CRISPR/Cas9 genome editing. The same embryos also expressed a ubiquitously expressed mCherry-tagged histone transgene to allow for cell lineage tracing. We traced cells from soon after fertilization (4-8 cell stage) through the last round of cell divisions for most cells (bean stage) by using StarryNite automated cell tracking software, and quantified reporter expression in each cell across time(25–28). Note, each Hox::GFP fusion line is homozygous viable, fertile, and displays no obvious phenotypes, demonstrating that the fusion proteins largely function like the wild type proteins.

**Figure 1:**
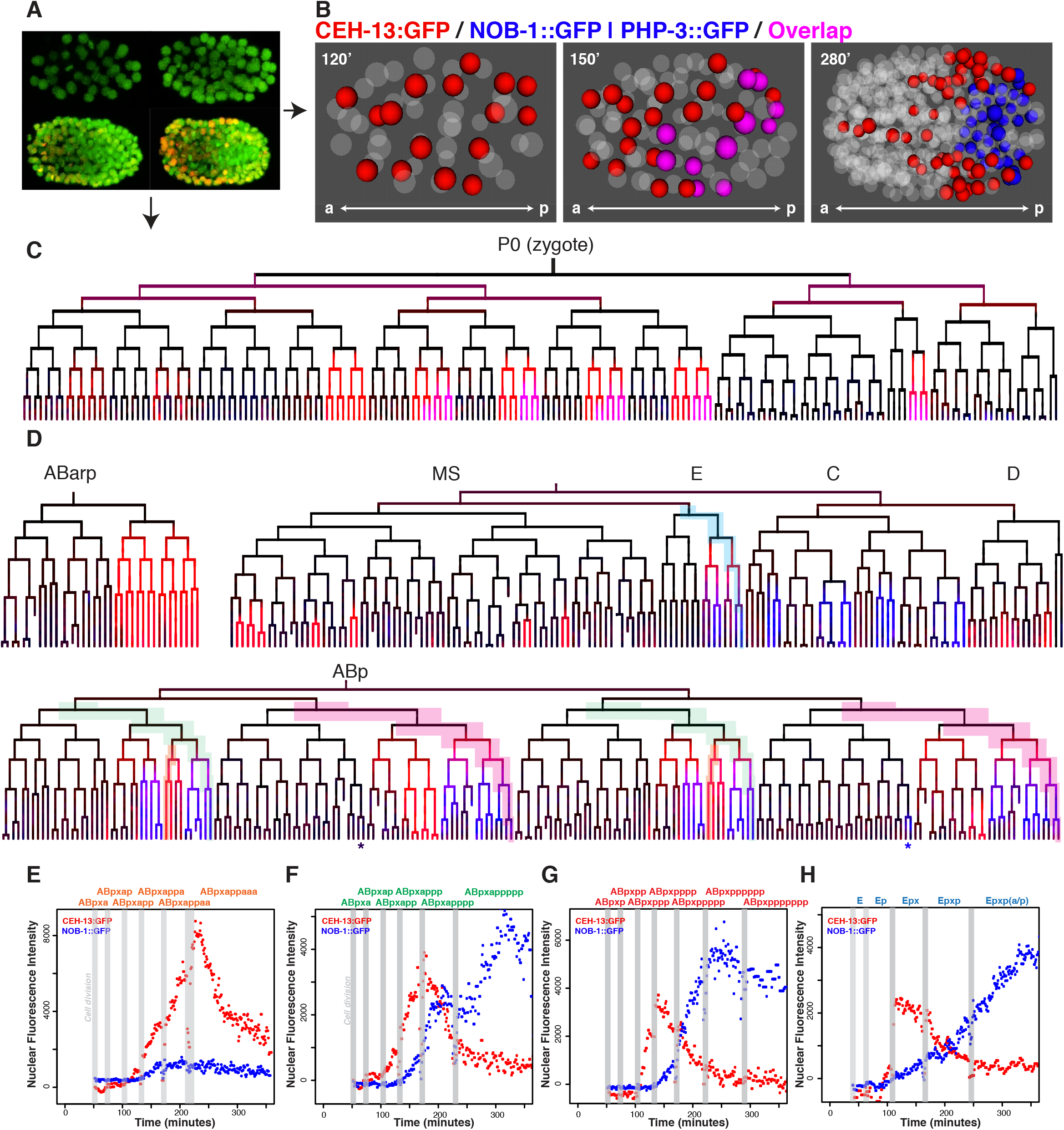
*ceh-13* and *nob-1/php-3* are expressed broadly in an overlapping pattern in the early *C. elegans* embryo. A) Time lapse images of transgenic *C. elegans* embryos carrying two transgenes, a ubiquitous fluorescent histone to mark all nuclei (shown in green) and a reporter of interest (shown in red), which can be a cis-regulatory element driving a fluorescent histone (transcriptional) or a fluorescently tagged transcription factor protein (translational). Image analysis software identifies nuclei, and quantifies reporter intensity within the nuclei, which can be displayed as a lineage tree colored by expression as in (C). B) 3D projections of nuclei expressing a CEH-13::GFP CRISPR reporter (red), NOB-1::GFP or PHP-3::GFP CRISPR reporter (blue), with overlap shown in magenta, at 120 minutes (50 cell stage), 150 minutes (100 cell stage), and 280 minutes (400 cell stage) post fertilization. NOB-1::GFP and PHP- 3::GFP CRISPR reporters were expressed in indistinguishable patterns (Supplemental Figure 1), but PHP-3::GFP was slightly brighter so is shown in the following panels. C) Lineage tree though the 100-cell stage, showing early expression of CEH-13 and PHP-3 CRISPR reporters, colored as in (B). D) Expressing lineages showing CEH-13 and/or PHP-3::GFP CRISPR reporter expression to 350 minutes of development, colors as in (B). Note that CEH-13::GFP precedes PHP-3::GFP and NOB-1::GFP in all lineages except ABp(l/r)papppp (asterisk - expression is consistent in ABprpapppp and variable in ABplpapppp). (E-H) Quantitative detail for highlighted lineages, showing nuclear fluorescence intensity of CEH-13::GFP (fosmid) and NOB-1::GFP transgene reporters across developmental time for the cells leading to ABp(l/r)appaaa (E), ABp(l/r)appppp (F), ABp(l/r)ppppppp (G), and Ep(l/r)p (H). For cell labels, x = (l/r). Nuclear fluorescence intensity is in arbitrary units. Grey bars mark cell divisions.

Consistent with previous studies using shorter reporters (20), CEH-13 expression is first seen eight Wnt-signaled cells (that are the more-posterior daughter after cell division) and their progeny at the 26-cell stage, during early gastrulation (Figure 1B,C, Supplemental Figure 2). These include the AB lineage-derived cells ABalap, ABalpp, ABarpp, ABplap, ABplpp, ABprap and ABprpp, and the posterior endoderm progenitor Ep. The ABalap and ABalpp expression was barely detectable and quickly faded, while the other lineages had more robust and persistent expression. . By the 200-350 cell stage, many of the granddaughters of the initially expressing cells have lost CEH-13 expression, while a few cells fated to become blast cells or neurons sustained or increased their expression. Also at this stage, some cells in the D and MS lineages as well as a few additional AB-derived cells activate CEH-13 expression (Figure 1D).

We compared the reporter expression pattern to that of endogenous mRNA as measured in a lineage-resolved single cell RNA-seq dataset, and found that the mRNA and reporter expression patterns were consistent (29). In comparison, a rescuing 35 kb CEH-13::GFP fosmid transgene had brighter expression in the cells expressing the CRISPR-tagged protein, plus additional weak expression detectable in some epidermal precursors from the C lineage (possibly detectable due to higher copy number). A shorter reporter containing only 8.2 kb of upstream sequence lacked expression in the MS lineage, emphasizing the role of distal regulatory elements in regulating *C. elegans* Hox gene expression (Supplemental Figure 1D) (30, 31).

Next, we defined the embryonic expression patterns of the posterior Hox genes *php-3* and *nob-1* tagged with GFP at the endogenous locus. As expected, *nob-1* and *php-3,* which are adjacent in the genome with *php-3* directly downstream of *nob-1*, and are expressed in identical patterns in both our GFP imaging data (Supplemental Figure 1A) and in the single-cell RNA-seq data (29). NOB-1/PHP-3 expression begins during mid-gastrulation in cells derived from the cells ABplapp, ABplppp, ABprapp, ABprppp, Ep, and the four posterior great-granddaughters of the C blastomere. Expression becomes stronger one cell cycle later in the AB-derived lineages with the exception of ABp(l/r)appa. Additional late embryonic expression occurs in the ABp(l/r)papppp lineages starting at the onset of morphogenesis (“bean” stage). NOB-1- expressing lineages give rise to fates including neurons, hypodermis, seam cells, epithelial cells, intestine, and death, and have posterior positions clustered in and near the developing tail at comma stage (32). Like CEH-13, early NOB-1::GFP expression persists through the terminal cell divisions in only a subset of cells (Figure 1D). A previously described NOB-1::GFP rescuing transgene that includes 9kb of sequence upstream of the *nob-1* transcript expresses GFP in a similar pattern, except it is not expressed in the C great-granddaughter lineages, suggesting this expression requires additional regulatory sequences outside of this region (Figure 1D, Supplemental Figure 1)(22).

Early CEH-13::GFP expression occurs in cells that are distributed broadly along the anterior-posterior axis (Figure 1B), while NOB-1 and PHP-3 expression is more limited to the posterior of the embryo. In the AB lineage, NOB-1/PHP-3 are expressed exclusively in cells that were the posterior sister after cell division and whose mother expressed CEH-13 (Figure 1D-H). In addition, these three genes are co-expressed at similar onset times in the posterior intestine lineage derived from Ep. In contrast, in late embryos, high levels of CEH-13::GFP and NOB- 1::GFP/PHP-3::GFP are largely mutually exclusive (Figure 1D, Supplemental Figure 1F, G), consistent with classic models of Hox expression and posterior dominance.

### Several overlapping lineage specific enhancers regulate *nob-1* embryonic expression

To identify regulators of *nob-1* embryonic expression, we tested nearby genomic sequences for embryonic *cis-*regulatory activity (Figure 2A,B). Previous work showed that other *C. elegans* Hox genes are regulated by distal enhancers located as much as 20kb from a given gene’s promoter (24,30,33), but enhancers for *nob-1* and *php-3* have not been identified. We took advantage of existing reporters of different lengths to narrow the sequence search space for *nob-1* embryonic enhancers (Figure 2C). A transcriptional reporter containing just 5.3 kb of upstream sequence driving histone-mCherry reporter expression, and the rescuing 9kb NOB- 1::GFP reporter are expressed in most of the same lineages as endogenous NOB-1::GFP, indicating they contain regulatory sequences sufficient for expression in full set of expressing lineages. However, while the protein fusion reporters are expressed at similar levels in each lineage, the shorter 5.3kb transcriptional reporter has much lower expression in the ABp(l/r)ppp lineages compared to other expressing lineages (Figure 2C, D). This suggests that additional sequences between -5.3kb and -9kb are required for the full endogenous expression levels in ABp(l/r)ppp.

**Figure 2:**
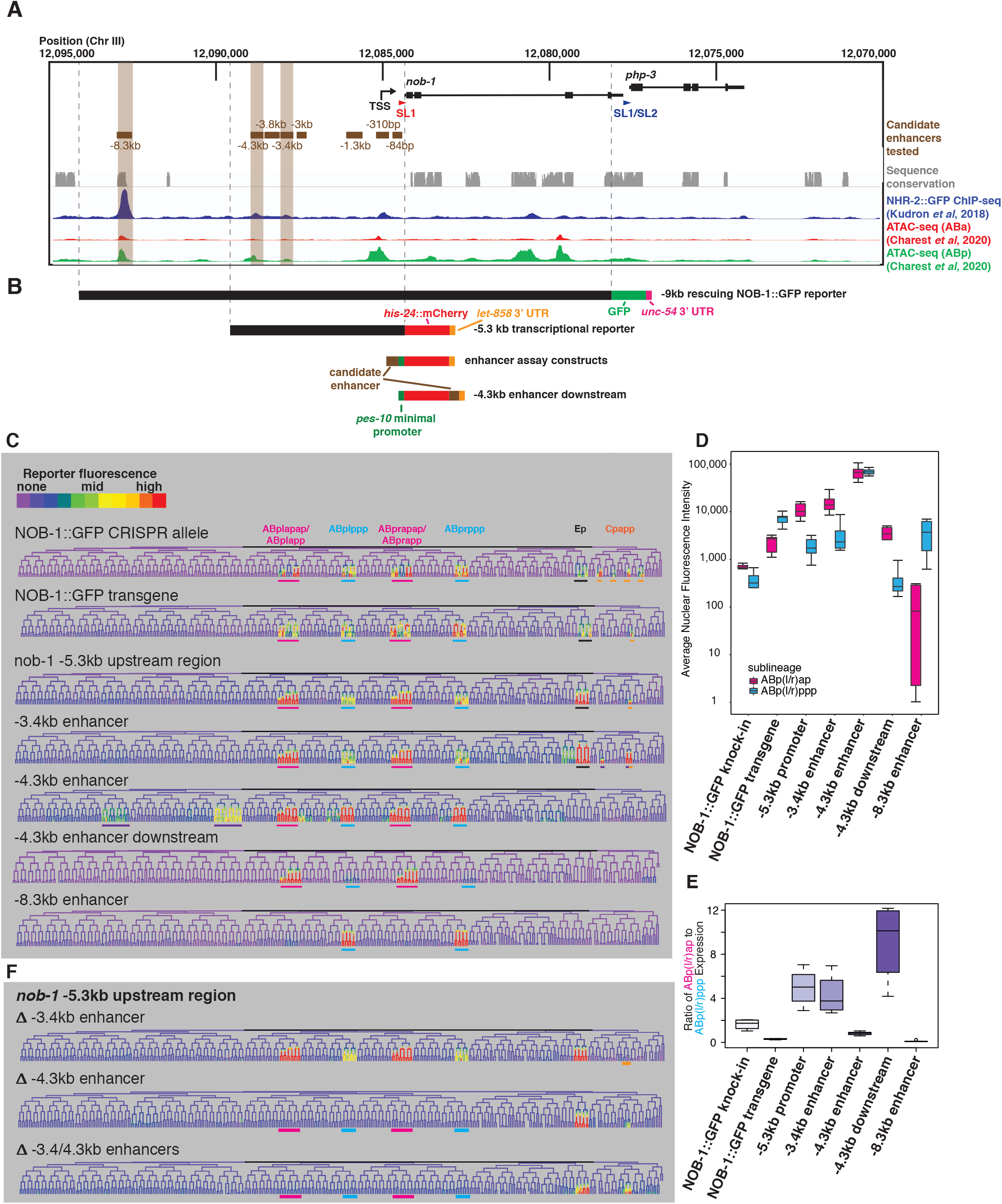
Regions upstream of *nob-1/php-3* can recapitulate its expression pattern. A) Genome browser view of the *nob-1*/*php-3* locus showing the genes (black), candidate enhancers tested (brown), sequence conservation with other nematodes (grey), and NHR-2 ChIP-seq trace (dark blue) from modENCODE (42) and ATAC-seq traces (red:ABa lineage, green: ABp lineage) from Charest et al, 2020 (65). B) Schematics of the *nob-1* reporter constructs examined, shown in alignment with (A). Enhancers were primarily tested in an orientation 5’ to the *pes-10* minimal promoter, but a downstream orientation was also used for the -4.3 kb enhancer. C) Lineage trees colored to show expression patterns for the various reporters and tested enhancers with relevant reproducible activity, using a rainbow color scale to increase visible dynamic range. Major expressing lineages are underlined: ABp(l/r)ap: pink, ABp(l/r)ppp: cyan, Ep: black, Cpapp: orange, ectopic: purple. Note the changes in expression driven by the -4.3 kb enhancer depending on its position relative to the *pes-10* minimal promoter. D) Average nuclear fluorescence values for cells in the ABp(l/r)ap (pink) and ABp(l/r)ppp (blue) lineages from at least four embryos, shown on a log scale. E) Ratio of expression in average ABp(l/r)ap cell to the average ABp(l/r)ppp cell at the 350 cell stage for at least 4 embryos. F) Lineage trees showing expression driven by the nob-1 -5.3 kb upstream region reporter with the -3.4 and/or -4.3 enhancers are deleted. Lineages where expression is lost are underlined with colors as in (C).

Although other *Caenorhabditis* species have large (12-20kb) intergenic regions upstream of *nob-1*, there is no detectable sequence conservation in the 5.3kb region upstream of *nob-1*, and only two short conserved stretches (85 and 275 nucleotides long) between 5.3 and 9kb upstream (34–37). This indicates that any conserved regulatory elements in this region have diverged substantially at the sequence level during *Caenorhabditis* evolution, limiting utility evolutionary conservation to identify enhancers. Chromatin marks typically used to identify transcriptional enhancers such as H3K27ac, H3K4me, or chromatin accessibility from whole embryos did not show strong peaks near *nob-1* or near 20 additional genes expressed in lineage-specific patterns at the same stage as *nob-1* (31,38–40).

We hypothesized that these chromatin signals are diminished in early embryonically expressed genes because the experiments used embryos of mixed stages where later stage nuclei dramatically outnumber nuclei from earlier stages. Therefore, we searched for other factors with published genomic binding patterns that could mark embryonic enhancers. One pattern stood out as preferentially bound near genes expressed with lineage-specific patterns in the early embryo: binding of a NHR-2::GFP fusion protein (41, 42). NHR-2::GFP binds in the intergenic sequences upstream of 19 of these 20 genes, and 16 genes have multiple clustered NHR-2 binding sites (vs. 22% and 8% of randomly chosen genes, respectively, p < 0.001; chi-squared test). NHR-2 is a nuclear hormone receptor distantly related to mammalian thyroid and PPAR receptors, and NHR-2::GFP is expressed in most or all somatic cells from the ∼50-cell to ∼200-cell stages. The functional importance of NHR-2 binding is unclear; *nhr-2* RNAi causes embryonic and larval arrest, but partial deletion alleles are viable. However, regardless of its function, we hypothesized that these clustered NHR-2 binding sites might be useful proxies for accessible chromatin and could predict enhancer activity in the early embryo.

We tested four NHR-2::GFP-bound regions upstream of *nob-1* for enhancer activity by generating transgenic worms expressing histone-mCherry under the control of each candidate enhancer placed upstream of a *pes-10* minimal promoter, which drives no consistent embryonic expression on its own (Figure 2A, B). We also tested four additional sequences for which we observed no NHR-2::GFP binding but which contained putative binding motifs for POP-1/TCF, which is required for expression of the *nob-1* transcriptional reporter (23). We identified embryonic cells expressing each enhancer reporter by confocal time lapse imaging and StarryNite. Only three of the regions (-3.4kb, -4.3kb and -8.3kb) tested showed enhancer activity that was consistent between embryos and also overlapped with the endogenous NOB-1::GFP expression pattern, suggesting they represent functional enhancers (Figure 2C, Supplemental Figure 2). All three functional enhancers were identified on the basis of NHR-2 binding.

The -3.4kb enhancer recapitulates most of the *nob-1* early embryonic expression pattern, with some differences in the C lineage, and drives much weaker expression in ABp(l/r)ppp than in ABp(l/r)app, similar to the *nob-1* -5.3 kb upstream transcriptional reporter. The -4.3kb enhancer also drives expression in ABp(l/r)app and ABp(l/r)ppp at a very high level (Figure 2D) and also variable misexpression in cells that don’t normally express NOB-1 (Figure 2C, Supplemental Figure 2A). When this enhancer was cloned downstream of the reporter, it showed less ectopic expression and stronger expression in ABp(l/r)app relative to ABp(l/r)ppp, similar to the -3.4 kb enhancer. This confirms the -4.3kb region as a *bona fide* enhancer capable of acting at a distance and suggests that the placement of this enhancer relative to the promoter influences its activity differently in different lineages and may be important for specificity. Embryos carrying multiple copies of the -4.3kb reporter transgene occasionally displayed the “no backend” phenotype observed in *nob-1/php-3* mutant embryos (Supplemental Figure 2B) (5). A third *nob-1* enhancer (-8.3kb) drives early embryonic expression only in ABp(l/r)ppp, as well as later expression in ABp(l/r)papppp. This most-distal enhancer is included in -9kb NOB- 1::GFP transgene but not the shorter -5.3kb transcriptional reporter, and includes the only substantial stretch of conserved sequence in the *nob-1* promoter region. This region thus can explain why ABp(l/r)ppp expression is relatively stronger in the translational reporter compared to the shorter transcriptional reporter.

As the -5.3kb *nob-1* transcriptional reporter includes both the -3.4kb and -4.3kb enhancers, we deleted each enhancer from this construct to test their necessity for *nob-1* expression (Figure 2C). Deleting both enhancers led to a complete loss of early embryonic expression in both AB lineages, suggesting there are no additional enhancers present sufficient for early embryonic expression in these lineages. The deletion of the -3.4 kb enhancer did not disrupt expression in ABp(l/r)app, ABp(l/r)ppp and in the absence of the -4.3 kb enhancer, this enhancer was not sufficient to rescue expression in these lineages. This indicates that while the -3.4kb enhancer may be sufficient to drive expression when placed directly next to the promoter, it is unable to drive expression in the endogenous context. Conversely, the -4.3 kb enhancer is a key driver of expression in these lineages. The expression remaining when both enhancers are lost indicates that additional sequences that drive the expression in the Ep and Cpapp lineages must exist within the -5.3 kb upstream region.

### *ceh-13* and Hox cofactors activate *nob-1* expression

Because *ceh-13* expression precedes that of *nob-1* in many lineages, we asked whether *ceh-13* is required for *nob-1* expression. To evaluate this, we examined *nob-1* reporter expression in embryos homozygous for the likely null mutation *ceh-13(sw1)* (21). Expression of the -5.3kb *nob-1* transcriptional reporter is eliminated in ABp(l/r)ppp (p < 0.001), and decreased by 25% in ABp(l/r)ap, and 40% in Ep (p < 0.02) (Figure 3A-C). Depletion of *ceh-13* by RNAi gave similar results with the transcriptional reporter (p < 0.002). In contrast, *ceh-13* RNAi does not significantly alter expression of the -9kb NOB-1::GFP transgene, indicating that there are additional cis-regulatory elements in this reporter which are *ceh-13* independent (Supplemental Figure 3A).

**Figure 3:**
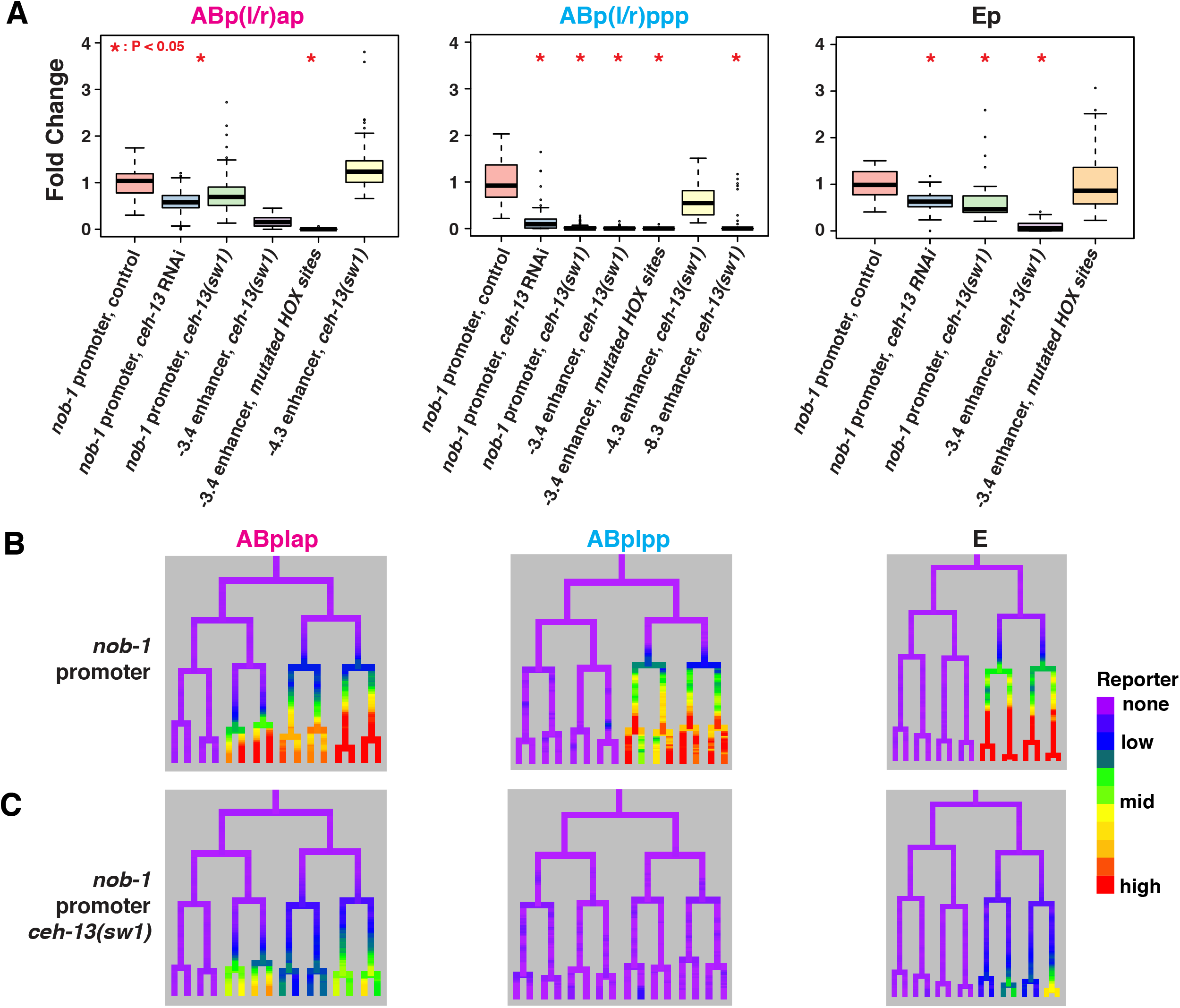
Expression of cis-regulatory elements upstream of *nob-1* and *php-3* depends on *ceh-13*. A) Fold change in expression level relative to mean wild-type control for *nob-1* promoter and enhancers in *ceh-13* mutant and RNAi conditions. *P* values determined by Wilcoxon Ranked Sum Test. B) Lineage view of wild-type *nob-1* -5.3kb upstream reporter (“promoter”) expression in specified lineages. C) -5.3kb *nob-1* promoter reporter expression in *ceh-13(sw1)* null mutant.

To determine which enhancers require *ceh-13*, we measured the activity of individual *nob-1* enhancer reporters in worms lacking *ceh-13* (Figure 3A, Supplemental Figure 3A). In *ceh- 13* mutants, expression driven by the -3.4 kb enhancer decreased significantly in all expressing cells, excluding the C lineage (p < 0.003) with an average decrease of 90% and complete loss in ABp(l/r)ppp. *ceh-13* loss also reduces activity of the -4.3 kb enhancer in ABp(l/r)ppp (47% decrease) but expression in ABp(l/r)ap is unchanged. This indicates that other factors besides *ceh-13* activate this enhancer in ABp(l/r)ap and suggests that this enhancer may be responsible for the residual expression of the 5.3kb transcriptional regulation in this lineage in *ceh-13* mutants. The distal -8.3kb enhancer also lost expression in ABp(l/r)ppp in the ceh-13(sw1) mutant (p < 0.002), indicating its activity also requires *ceh-13*. Motif analysis identified six putative *ceh-13*-binding motifs in the -3.4kb enhancer; mutating these sites abolished expression in the AB and C lineage (p < 0.02, p < 0.03), but did not alter expression in the Ep lineage indicating *ceh-13* regulation of *nob-1* may be indirect in the E lineage (Figure 3A, Supplemental Figure 3B). These results show that multiple *ceh-13*-dependent enhancers work together to regulate *nob-1* expression in gastrulating embryos.

Expression of classically defined Hox targets often require Hox cofactors such as *homothorax* or *extradenticle*. These cofactors form larger TF complexes with Hox factors to increase binding specificity and may also have Hox-independent functions (43–45). To determine when and where Hox cofactors are expressed in early embryos, we used StarryNite to trace the expression of translational reporters for *unc-62*, the *C. elegans homothorax/Meis* homolog, and of *ceh-20* and *ceh-40*, the homologs of *extradenticle/Pbx*. A third *extradenticle* homolog, *ceh-60*, is only expressed in later development (>200 minutes) and does not overlap with *ceh-13*, so was not investigated further(29). We found that each of these genes has specific and dynamic expression patterns that overlap with each other and with *ceh-13* and *nob- 1* expression (Figure 4A-D, Supplemental Figure 5). In particular, all three cofactors are coexpressed with CEH-13 in cells that will later express NOB-1. Other CEH-13 and NOB-1 expressing cells also express all three cofactors, except only CEH-20 is expressed in the early E lineage, and all three are lost in the NOB-1-expressing ABp(l/r)appp lineage as the embryo approaches morphogenesis. We conclude that Hox cofactors are expressed in cells where *ceh- 13* activates *nob-1* expression.

**Figure 4:**
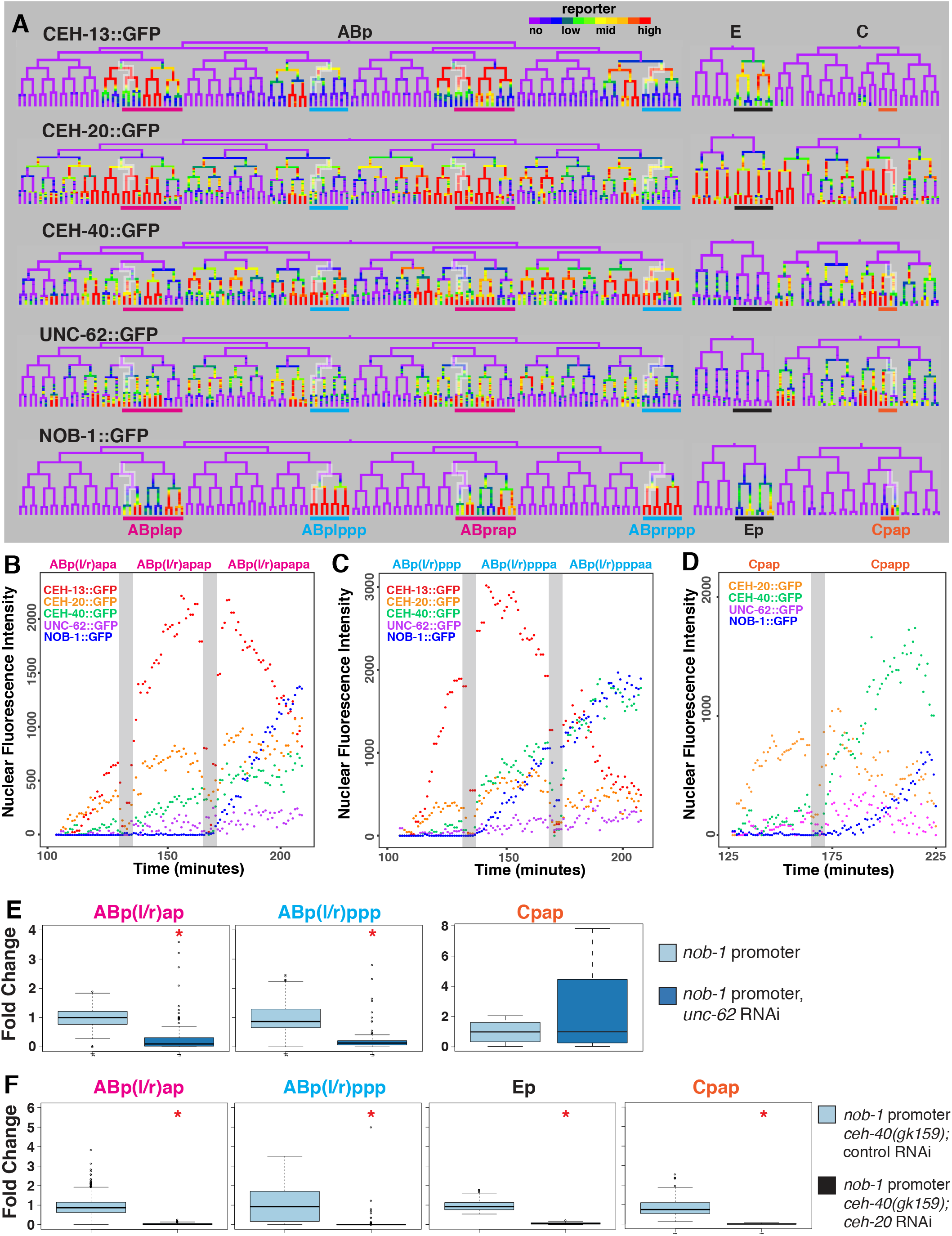
Hox co-factors precede *nob-1* and regulate its expression. A) Trees of NOB- 1::GFP expressing lineages, ABp, E, and C, showing the expression of *ceh-13* (fosmid) and *nob-1* (GFP transgene) reporters, and fosmid GFP transgene reporters for the Hox co-factors, *ceh-20*, *ceh-40* and *unc-62*. Color thresholds were adjusted for each gene to show all expressing cells. Highlighted branches are shown in graphs B, C and D. B-D) Average TF reporter nuclear fluorescence intensity across embryos (n >= 2) and for left/right symmetric cells across developmental time for the cells leading to B) ABp(l/r)apapa, C) ABp(l/r)pppaa and D) Cpapp. Fluorescence intensity is in arbitrary units and grey bars mark cell divisions. E,F) Fold change values for -5.3 kb *nob-1* promoter expression in untreated and *unc-62* RNAi treated (E) or *ceh-40(gk159)* mutant embryos treated with *ges-1* (control) or *ceh-20* RNAi (F) in specified lineages. Significant changes marked by asterisk; *P* values determined by Wilcoxon Ranked Sum Test.

We tested whether Hox cofactors regulate *nob-1* expression by examining *nob-1* reporter expression after loss of each gene. We found that depleting *unc-62* by RNAi led to significant loss of expression of the *nob-1* -5.3kb reporter in all AB lineages (80% decrease, p=0.001), with near complete loss in the ABp(l/r)apap and ABp(l/r)ppp lineages (Figure 4E). The -9kb NOB-1:GFP protein reporter showed similar expression decreases, although this was only significant in ABp(l/r)appa and incompletely penetrant in other lineages (Supplemental Figure 4B-C). In contrast, CEH-13::GFP reporter expression was unchanged after *unc-62* RNAi (Supplemental Figure 4D). These results show that *nob-1* is regulated by *unc-62*. The fact that *unc-62* RNAi has a stronger phenotype in ABp(l/r)appa than loss of *ceh-13* suggests that *unc-62* may regulate *nob-1* independently of *ceh-13* in these cells.

To investigate the role of the *extradenticle* homologs, we examined the *nob-1* transcriptional reporter in a *ceh-40(gk159)* null mutant background in embryos of worms treated with control or *ceh-20* RNAi. We found that the combination of *ceh-20* RNAi with the *ceh-40* mutation caused a significant loss of expression in virtually all expressing cells including Ep and Cpap as compared to control RNAi (97% decrease, p < 0.004). This indicates that the *exd* homologs *ceh-20* and *ceh-40* are required to activate *nob-1* expression and that they are at least partially independent of *ceh-13* in the ABp(l/r)ap, Ep, and Cpap sublineages. This finding is consistent with previous reports that *nob-1* RNAi causes increased lethality in a *ceh-40* mutant background relative to wildtype animals (46).

### Multiple lineage-specific embryonic TFs including *elt-1/GATA*, *ceh-36/OTX* and *unc- 30/PITX* are required for early *nob-1* expression

Since some NOB-1::GFP reporter expression remains in the absence of *ceh-13*, especially in the ABp(l/r)ap lineage, other factors must also activate *nob-1* in these cells. Indeed, we previously identified the Wnt effectors *pop-1* and *sys-1* as activators of *nob-1* expression (23). We took advantage of databases of TF expression to search for additional candidate regulators (29,40,47).

Only two of the three enhancers (-3.4kb and -4.3kb) drive expression in the ABp(l/r)ap lineage, so to identify additional potential regulators of *nob-1* in this lineage, we used two criteria. First, we searched for TFs expressed specifically in ABp(l/r)ap, but not ABp(l/r)pp. Second, we used a TF binding site scanning approach to identify which of these TFs have predicted binding motifs in the -3.4kb and -4.3kb enhancers (9, 48). From this analysis, the GATA zinc finger encoded by *elt-1* emerged as the strongest candidate. An ELT-1::GFP reporter is expressed prior to gastrulation in the ABp(l/r)ap progenitors ABpla and ABpra, in the other major epidermis-producing lineages ABarp, Caa and Cpa, and in the primarily neuronal lineages ABalap and ABalppp (Figure 5A). ELT-1::GFP expression is later downregulated in cells that do not adopt epidermal fates (Figure 5B). ELT-1::GFP expression in the ABpla/ABpra lineages could be detected at least 20 minutes prior to *nob-1* reporter expression in ABp(l/r)ap (Figure 5B, C). GATA Motifs predicted to bind ELT-1 are enriched in the ABp(l/r)ap enhancers - 3.4kb (8 sites) and -4.3kb (7 sites) enhancers compared to the -8.3 enhancer (1 site), which does not drive expression in ABp(l/r)ap. While the *C. elegans* genome encodes several other GATA factors, only *elt-1* is expressed prior to *nob-1* in the ABp(l/r)a linage(29, 49).

**Figure 5:**
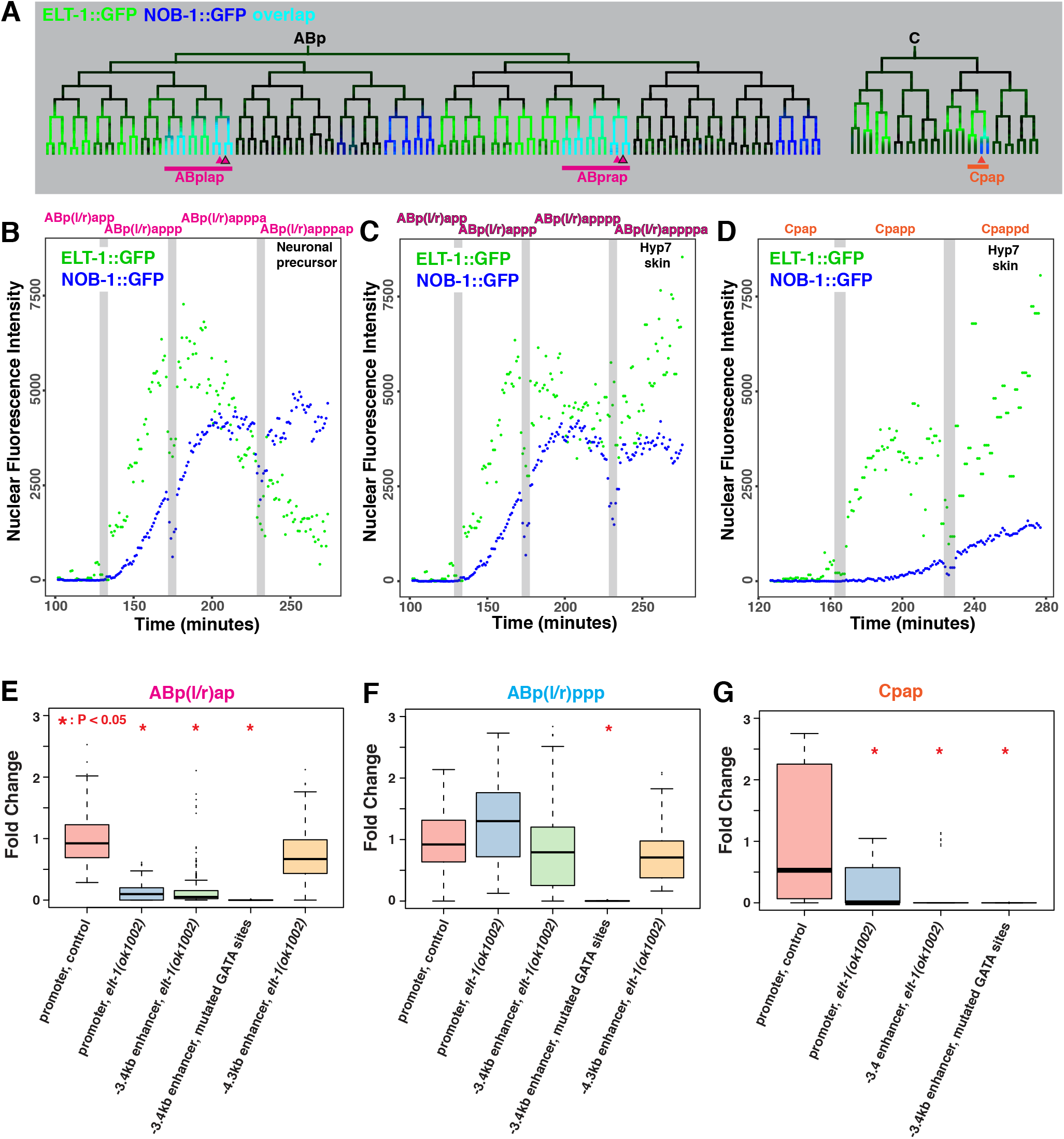
*elt-1* regulates *nob-1* expression in ABp(l/r)ap lineage. A) Partial lineage trees showing ELT-1::GFP (Green), NOB-1::GFP (Blue) or overlapping (Cyan) expression in the ABp and C lineages. Arrowheads indicate branches shown in subsequent panels. B-D) Average ELT-1::GFP and NOB-1::GFP nuclear fluorescence intensity (arbitrary units) in cells leading to ABp(l/r)apppap (neural fate) C), ABp(l/r)appppa (hypodermal fate), and D) Cpappd (hypodermal fate). Grey bars mark cell divisions. E) Fold change values for the *nob-1* promoter and -3.4 kb and -4.3 kb wild-type or mutated enhancers in the ABp(l/r)ap lineage in the control and *elt-1(ok1002)* mutant conditions. F-G) Fold-change of *nob-1* reporter intensity in *elt- 1(ok1002)* relative to mean wild-type control level in the E) ABp(l/r)pp, F) ABp(l/r)ppp, and G) Cpap lineages. *P* values determined by Wilcoxon Ranked Sum Test.

To test whether *elt-1* regulates *nob-1* expression, we measured the activity of *nob-1* reporters in the *elt-1(ok1002)* null mutant background (Figure 5E-G, Supplemental Figure 6A,B). We found that *elt-1* mutants showed decreased expression of the *nob-1* transcriptional reporter in the ABp(l/r)ap lineage (87% decrease, p < 0.014), and the Cpap lineage (92% decrease, not significant), in which *elt-1* precedes *nob-1* expression, but not in ABp(l/r)pp, where ELT-1::GFP is not expressed. Similarly, we observed decreases in the same sublineages for the -3.4kb enhancer reporter (ABp(l/r)ap: 72% decrease, p < 0.008; C: 71% decrease, p < 0.03) and the - 4.3kb enhancer reporter (ABp(l/r)ap: 31% decrease, not significant). To determine if *elt-1* directly regulates the -3.4kb enhancer, we mutated the predicted ELT-1 binding sites, which have a characteristic GATA motif, and observed loss of reporter expression in all lineages (p < 0.001). We conclude that *elt-1* activates *nob-1* expression in the ABp(l/r)ap lineage through at least two enhancers, and is at least partially redundant with other factors that may bind GATA motifs.

We previously showed that two homeodomain *bicoid* family TFs, *ceh-36/OTX* and *unc- 30/PITX* are redundantly required for proper development of the ABp(l/r)p lineage (50). Since *ceh-13* and *elt-1* mutants still have some *nob-1* reporter expression in the ABp(l/r)p descendants, we hypothesized that *ceh-36* and *unc-30* might be important for *nob-1* expression in these lineages. All three enhancers contain at least two putative CEH-36/UNC-30 binding sites, and the -8.3 kb enhancer, which activates expression only in ABp(l/r)p, has nine (Supplemental Tables S3-S5). To test this hypothesis, we examined NOB-1::GFP transgene expression in eight *ceh-36;unc-30* double mutant embryos, and found that two of the eight embryos (25%) lost expression in this lineage (Supplemental Figure 7A). Furthermore, the ratio of ABp(l/r)ap to ABp(l/r)ppp expression significantly increased (p < 0.002) in the remaining double mutant embryos, consistent with a decrease in ABp(l/r)ppp expression (Supplemental Figure 7B). In conclusion, at least seven partially redundant lineage-specific transcription factors positively regulate *nob-1* expression: *ceh-13, ceh-20, ceh-40, unc-62, elt-1*, *ceh-36*, and *unc-30*.

### ceh-13 and nob-1 are required for correct cell position and division patterns in the early embryo

Previous work suggested that *nob-1/php-3* and *ceh-13* are required for cell position but not other aspects of cell fate specification (5). However, mutating lineage identity regulators often leads to defects in cell cycle length or division patterns (50, 51) that could have been missed in previous studies. To test for such defects, we measured cell positions and cell division timing in six *nob-1/php-3* (referred to as nob-1 for simplicity) and *ceh-13* mutant embryos by time-lapse microscopy and StarryNite cell tracking, and compared them to a database of 17 wild type embryos (27, 50). (Figure 6)

**Figure 6:**
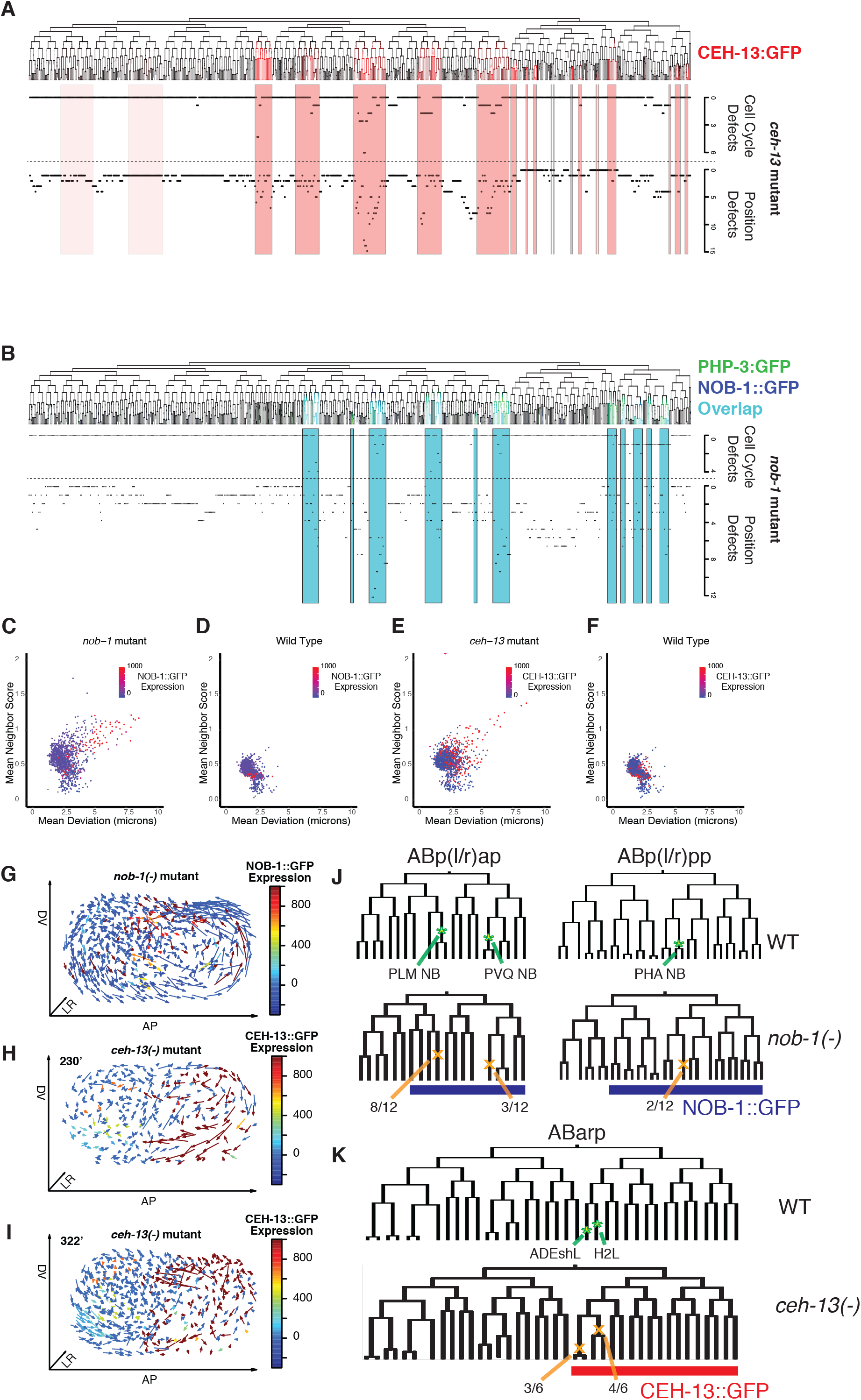
Cell defects are observed in the posterior of both *ceh-13* and *nob-1* mutant embryos. A) Lineage showing *ceh-13* expression with plots of frequency of cell cycle defects and cell position defects for each cell plotted below. Cells expressing *ceh-13* during their development are highlighted in pink (ABalap and ABalpp, which express very low levels of CEH- 13::GFP, are denoted with lighter shading). Cell cycle defects are defined as missed, ectopic or significant change in division timing of at least 5 minutes and cell position defects are defined as significant deviation of at least 5 μm from the positions expected from wild type (see methods). Defects are listed cumulatively for terminal cells, so defects in both a parent and daughter cell would be scored as two defects. B) Lineage showing *nob-1* and *php-3* expression overlap with plots of frequency of cell cycle defects and cell position defects for each cell plotted below. Cells expressing both *nob-1* and *php-3* are highlighted in cyan on the plots. C, D) Plot of mean deviation in microns vs. mean neighbor score for each cell in the embryo, colored by level of NOB-1::GFP expression, for *nob-1(ct223)* mutant which disrupts both *nob-1* and *php-3* (C) and wild-type (D) embryos. Neighbor score is a measure of whether a cell is close to its normal neighbors; values above 0.8 indicate aberrant neighbor relationships. E, F) Plot of mean deviation in microns vs. mean neighbor score for each cell in the embryo, colored by level of CEH-13::GFP expression, for ceh-13(sw1) mutant (E) and wild-type (F) embryos. (G-I) Three-dimensional plot of cell position deviations - arrows point from average wild-type location to average mutant location for each cell, colors indicate the level of NOB-1::GFP (G) or CEH- 13::GFP(H-I) expression: G) *nob-1(ct223)* at 335 minutes of development. H) *ceh-13(sw1)* at 230 minutes of development) *ceh-13(sw1)* at 322 minutes of development. J) wild-type and *nob- 1(ct223)* mutant lineages for the ABp(l/r)ap and ABp(l/r)pp lineages, showing examples of cell division defects. Green stars indicate the normal divisions of neuroblasts, orange Xs indicate the failed divisions of these cells with the frequency observed noted below. Blue underline indicates the expression of NOB-1::GFP. K) Wild-type and *ceh-13(sw1)* mutant lineages for the ABarp lineage. Green stars indicate the normal development of ADEshL and H2L cells, orange Xs indicate the ectopic division of these cells with the frequency observed noted below. Red underline indicates the expression of CEH-13::GFP.

Consistent with previous work (5), we observed striking global patterns of cell position defects in *nob-1* mutant embryos. The most severe and highest penetrance defects were in cells from NOB-1-expressing lineages (Figure 6A,C,D,G, Supplemental Figure 8)). In total, 48% (62 of 127) of cells descended from NOB-1::GFP expressing lineages had severe position defects (>5 micron deviation, z score > 3.5) in at least two embryos compared to 1.2% (14 of 1094) of cells from non-expressing lineages (p <<10^-20^). However, a large number of cells from non-expressing lineages were moderately mis-positioned, suggesting that many defects are not cell-autonomous. Globally, dorsal cells that normally express NOB-1 fail to migrate to the posterior at 230 minutes post-fertilization; by 300 minutes non-expressing ventral cells compensate by inappropriately moving to the posterior, resulting in a global counterclockwise rotation of cell positions when the embryo is viewed from the left aspect (Figure 6G). The position defects we observed are consistent with the severe posterior morphology defects observed in rare surviving larvae.

We also identified broad cell position defects in *ceh-13* mutant embryos (Figure 6B,E,F,H,I). Intriguingly, the global patterns of cell position defects differed from *nob-1* mutants. The number of severe position defects was lower for *ceh-13* than for *nob-1*, with 4% (15/357) of cells descended from CEH-13::GFP expressing lineages severely displaced in at least two embryos, as opposed to 0.5% (5/874) of cells from non-expressing lineages (p << 10^-20^). At 210 minutes, (200 cell stage), the most strongly mispositioned cells were *ceh-13-*expressing cells that do not express *nob-1* in the posterior ventral region of the embryo. At this stage, cells on the dorsal surface were posteriorly mispositioned, such that cells in the posterior half of the embryo were rotationally mispositioned clockwise, the opposite direction to that seen in *nob-1* mutants (Figure 6H). Cells at the anterior ventral surface (both *ceh-13*-expressing and non-expressing) were mispositioned towards the posterior, resulting in a counterclockwise rotational defect in the anterior half of the embryo. At later stages, these global defects were largely resolved, and after the onset of morphogenesis the position defects were largely limited to a small number of strongly mispositioned cells (Figure 6I). The most extreme were the DA and SAB motorneurons after 300 minutes (bean stage), which were mispositioned several cell diameters anterior of their wild-type position (Supplemental Figure 8B). Given the recovery of most cell positions by the beginning of morphogenesis, it is unclear if cell position defects contribute to the severe anterior morphogenesis phenotypes in *ceh-13* mutants that hatch (21), although the earlier cell position defects could disrupt normal cell-cell signaling interactions required for fate specification.

In contrast to previous studies (5, 21), we also identified numerous defects in cell division timing or pattern in both *nob-1* and *ceh-13* mutants, and these were also heavily restricted to expressing lineages (Figure 6A,B). 13% of *nob-1* expressing cells and 5% of *ceh-13* expressing cells had severe division-time defects in at least two of the corresponding mutant embryos (>5 minute deviation from wild type, and z-score >5), vs none in non-expressing cells (p << 10^-20^). The most common cell division defect in *nob-1* mutants was the PLM/ALN neuroblast mother (ABp(l/r)apappp), which failed to divide in 8 of 12 observations (Figure 6J). The neuroblast that normally produces the PHA, PVC and LUA neurons instead underwent programmed cell death in two of 12 mutant lineages. Other cells that had multiple cell cycle delays in *nob-1* mutants included the PVQ neuroblast, PHB/HSN neuroblast, and the mother of the epidermal cells PHsh, hyp8 and hyp9. All of these cells also were mispositioned (mean 5 micron deviation from expected position, Supplemental Figure 8C).

The most striking defects in *ceh-13* mutants were in the cells ABarppaaa(a/p), which normally differentiate into the ADE sheath (glial) cell and H2 epidermal cell, but instead each divided inappropriately in 4 of 6 mutant embryos (Figure 6K). In addition, ABplppaaap, which normally undergoes programmed cell death, instead divided in two *ceh-13* mutant embryos. Several neuroblasts including those producing the DA, DD and SAB motorneurons also had cell division delays in *ceh-13* mutants. We conclude that both *ceh-13* and *nob-1/php-3* are required not only for proper cell positioning, but also for normal division patterns, suggesting a broader role in fate specification. Furthermore, the incomplete penetrance of both cell cycle and cell position defects in *ceh-13* and *nob-1* mutants indicate redundancy in developmental programming that likely contributes to the robustness of *C. elegans* embryonic development (Sulston, 1983).

## Discussion

Classic work defined the concept of Posterior Dominance, in which posterior Hox genes repress the expression and activity of more anterior Hox genes (1). Indeed, previous work in later developmental stages showed that *nob-1* represses *ceh-13* in the posterior ventral nerve cord, consistent with posterior dominance (45). In contrast, we show that *ceh-13* activates *nob- 1/php-3* expression through at least two enhancers during *C. elegans* gastrulation. To our knowledge, this is the first example of an anterior Hox gene positively regulating the expression of a posterior Hox gene during development. *ceh-13* and *nob-1/php-3* are also expressed at an earlier phase in development (pre-gastrulation for *ceh-13* and mid-gastrulation for *nob-1/php-3*) than in many other organisms. These observations, along with the lineage defects seen in *ceh- 13* and *nob-1* mutants, suggest that the role of *ceh-13* and *nob-1* is to regulate lineage identity, distinct from their later role in positional identity. The role of Hox cofactors such as *unc-62, ceh- 20* and *ceh-40* in early embryonic *nob-1* regulation suggests that these novel functions in lineage specification nonetheless share some common mechanisms with the later roles. The fact that *C. elegans* is largely unsegmented may have allowed the cooption of these Hox genes for this distinct role. This emphasizes the striking amount of flexibility in the *C. elegans* Hox cluster.

Several studies have highlighted that conserved transcription factors often “moonlight,” with the same factor playing apparently distinct roles in early progenitors and in terminal cell types. For example, the OTX homeodomain TF *ceh-36*, and the PITX homeodomain TF *unc-30*, which both specify specific terminal neuron types, separately act redundantly 6-7 cell divisions earlier in the ABp(l/r)p lineages to specify broad features of lineage identity (50,52–54), and *skn- 1*, which is required maternally to specify the endomesodermal blastomere EMS at the 4-cell stage is required postembryonically for oxidative stress resistance and longevity (55, 56). Our work shows that *ceh-13* and *nob-1/php-3* are another example of this phenomenon. Our identification of cell division defects in each mutant demonstrates they also play a role in lineage specification. In addition, we confirmed that each is also required for correct cell position; it is important to note that position defects could also result from incorrect specification of lineage identity. Previous work showed that at least one neuron (the PLM touch receptor) is defective in less severe *nob-1* mutants(19); the failure of PLM neuroblasts to divide in *nob-1/php-3* mutants indicates a function in PLM progenitor lineages. Similarly, *ceh-13* mutants have cell division defects, and *ceh-13* is required for the lineage-specific activity of specific *nob-1* enhancers. Finally, while *elt-1* has a well-defined known role in epidermal fate specification(57), our work shows that it is also required for *nob-1* enhancer activity. This is consistent with other recent work identifying defects in cell division patterns and the expression of the neurogenic TF *lin-32* in *elt-1* mutants(58). The ability of developmental transcription factors to “moonlight’ is likely facilitated by complex regulation by multiple enhancers as seen for *nob-1/php-3*, and *ceh- 13*(30, 31) since enhancer evolution can enable genes to be expressed at different locations and times in development(59, 60). An intriguing question is whether these multiple Hox functions are made possible by the unique nature of the *C. elegans* lineage, where positional identity is defined at least in part by lineage history (17, 61), or if they are conserved in other species.

While early reporter studies found that the expression of many genes expressed in terminal cells is well approximated using just the promoter-proximal region(62–64), open chromatin mapping studies have identified many distal regions of open chromatin, many of which can act as enhancers (31,38,39). Our work reinforces these studies and emphasizes the importance of enhancers in *C. elegans* gene regulation. Intriguingly, other genes expressed in lineage-specific patterns around the same time as *nob-1* have similar tendency to have multiple nearby regions of NHR-2::GFP binding and lineage-specific accessible chromatin as measured by ATAC-seq (65), suggesting they may also be regulated by multiple distal enhancers. This could reflect these genes’ need for more complex regulation. The *nob-1/php-3* locus contains at least two or three enhancers responsible for expression in each expression domain during gastrulation. This is reminiscent of so-called “shadow enhancers” identified in other organisms, for which redundancy appears to confer robustness in the face of environmental variability(66–68). This high level of redundancy also appears to extend to the number of transcription factors regulating each enhancer. Current data implicate at least eight TFs as activators of *nob-1* enhancers including *ceh-13*, *elt-1*, *ceh-36*, *unc-30*, the Wnt effector *pop-1*, and the Hox co- factors *ceh-20*, *ceh-40* and *unc-62* (Supplemental Figure 9). In some cases, the same binding motif may be regulated by different factors in different lineages. For example, mutating six GATA motifs in the -3.4 kb enhancer caused the loss of expression in ABp(l/r)ap, potentially due to loss of ELT-1 binding. However, this mutant also lost expression in Ep, which does not express *elt-1*, but does express other GATA factors with similar binding specificity including *end-1*, *end-3*,and *elt-7* (69). Similarly, the more severe loss of *nob-1* enhancer expression when CEH-13 motifs are mutated than observed in *ceh-13* mutant animals indicates that other TFs may also bind these sites. The partially penetrant nature of many of the mutant phenotypes further emphasizes the high level of redundancy. In sum, multiple modes of regulatory redundancy could provide a mechanism for the remarkable developmental robustness observed in *C. elegans*.

## Methods

### Strain generation and propagation

Worm strains (Supplemental Table 1) were maintained at 21-23°C on OP50 *E. coli* on NGM plates. Enhancer reporter strains were generated by injection into RW10029, a GFP histone strain used for lineage tracing. Injection cocktails consisted of reporter DNA construct at 10ng/μL, with 5 ng/μL myo-2p::GFP and 135 ng/μL pBluescript vector and were injected using a Narishige MN-151 micromanipulator with Tritech microinjector system. The *nob-1* promoter deletion strains were generated by injection and compared to a wildt-ype *nob-1* promoter strain created by injection of pJIM20::nob-1, JIM518. The *ceh-13* rescuing fosmid, *ujIs153*, was created by bombardment of WRM0622C_C06 (70) as previously described (50). Other strains were created through crosses using standard approaches. Lethal alleles were maintained as balanced heterozygotes (*ceh-13(sw1))* or rescued by free duplications (*nob-1(ct223)*), with homozygous mutants recognized by characteristic high-penetrance morphology defects. Note, the *ct223* allele disrupts both *nob-1* and *php-3*. Lethal *ceh-36;unc-30* double mutants were maintained as homozygotes rescued by an extra-chromosomal array carrying *ceh-36*::GFP; lack of rescue was scored by absence of GFP in the ABpxpa lineage. CEH-13::GFP and NOB- 1::GFP CRISPR knock-in alleles were created under contract by SUNY Biotech (Fuzhou, China).

### Molecular biology

Candidate enhancers were amplified with Phusion HF polymerase (New England Biosciences) from either pJIM20::nob-1 or N2 genomic DNA, with overhangs for stitching, which were then either gel or PCR purified (Qiagen). Putative enhancers were attached to a *pes-10* minimal promoter::HIS-24::mCherry::let-858 3’UTR fragment amplified from POPTOP plasmid (71) (Addgene #34848) by using PCR stitching to create an enhancer reporter which was sequence verified and purified with a PureLink PCR purification kit (ThermoFisher) for injection. The *pes- 10* minimal promoter drives no consistent early embryonic expression alone, but has previously been shown to facilitate expression driven by different enhancers (71–73). For the downstream enhancer experiment, the -4.3kb enhancer was cloned immediately downstream of the reporter stop codon. Tested enhancer sequences can be found in Supplemental Table 2. Regions deleted from the -5.3 kb transcriptional reporter match the tested enhancer sequences exactly. Putative transcription factor binding sites were identified using CIS-BP (cisbp.ccbr.utoronto.ca)(9, 48) (Supplemental Table 3-5). DNA fragments with desired mutations were synthesized (Integrated DNA Technologies) and PCR stitched as before.

### Imaging/Lineaging

We acquired confocal images with a Leica TCS SP5, Stellaris or Nikon A1RSi resonance scanning confocal microscope (67 z planes at 0.5 μm spacing and 1.5 minute time spacing, with laser power increasing by 4-fold through the embryo depth to account for attenuation of signal with depth). Embryos from self-fertilized hermaphrodites were mounted in egg buffer/methyl cellulose with 20μm beads as spacers (74) and imaged at 22°C using a stage temperature controller (Brook Industries, Lake Villa, IL). We used StarryNite software to automatically annotate nuclei and trace lineages, and AceTree software to identify and fix any errors from the automated analysis, and quantified reporter expression in each nucleus relative to local background (using the “blot” background correction technique) as previously described (25,26,28,75,76).

### Quantitative comparisons

Cell averages of nuclear fluorescence were computed for each cell based on all (typically >20) measurements across its lifetime (Supplemental Table 6). For each control condition, the mean control value of each named expressing cell was computed and used to determine the fold change for each control and mutant cell with the same name, which were displayed as boxplots using R 3.5.1 (The R Foundation for Statistical Computing). Background-corrected expression values were rounded up to zero if values fell below zero. To determine if changes were significant across the lineage, the summed expression for each cell in the lineage of interest was calculated for all control and mutant embryos and the two sets of values were compared using a Wilcoxon Ranked Sum test using R. For each condition, at least four control and three mutant embryos were analyzed (exact numbers in Supplemental Table 7). Each embryo analyzed represents an independent biological replicate. To evaluate the enrichment of NHR-2 binding sites, a chi-squared test was used.

### Mutant cell position analysis

Cell position defects were identified as previously (50). Briefly, we corrected for differences in global division rates (which did not differ dramatically from wild-type), and considered divisions as defective/outliers if they deviated from the wild-type cell cycle length by at least five minutes and had a z-score greater than three (Supplemental Tables 8-10). Cell positions (Supplemental Tables 11-13) were corrected for differences in embryo size and rotation, and considered defective if they deviated from the expected wildtype position by at least five microns, had a z- score greater than five, and a nearest neighbor score greater than 0.8 (defined empirically based on the distribution of wild-type scores).

## Acknowledgements

We thank Brian Gebelein and members of the Murray, Zacharias, and Gebelein laboratories and the Philly Worm Group for helpful discussion and comments on the manuscript. We thank Bob Waterston and Adrian Streit for strains. Some strains were provided by the CGC, which is funded by NIH Office of Research Infrastructure Programs (P40 OD010440). We thank WormBase, which is supported by NHGRI grant U24 HG002223, for consolidating valuable information. We thank Swati Mundre, Tom Sesterhenn,Yannis Belloucif, Ilona Jileaeva for cloning assistance and technical assistance with lineage tracing and worm genetics. We thank Meera Sundaram for the use of injection equipment and reagents. We thank Stephen Nehrbass for software assistance and Matt Kofron and Evan Meyer at the Confocal Imaging Core at Cincinnati Children’s, a Nikon Center of Excellence, for their help with imaging, and Matt Batie for designing and building the cooling stage. This work was supported by NIH grants R35GM127093 (to J.I.M.), R00GM111825 (to A.L.Z.), and F31GM123737 (to J.D.R.).

---

**Supplemental Figure 1:**
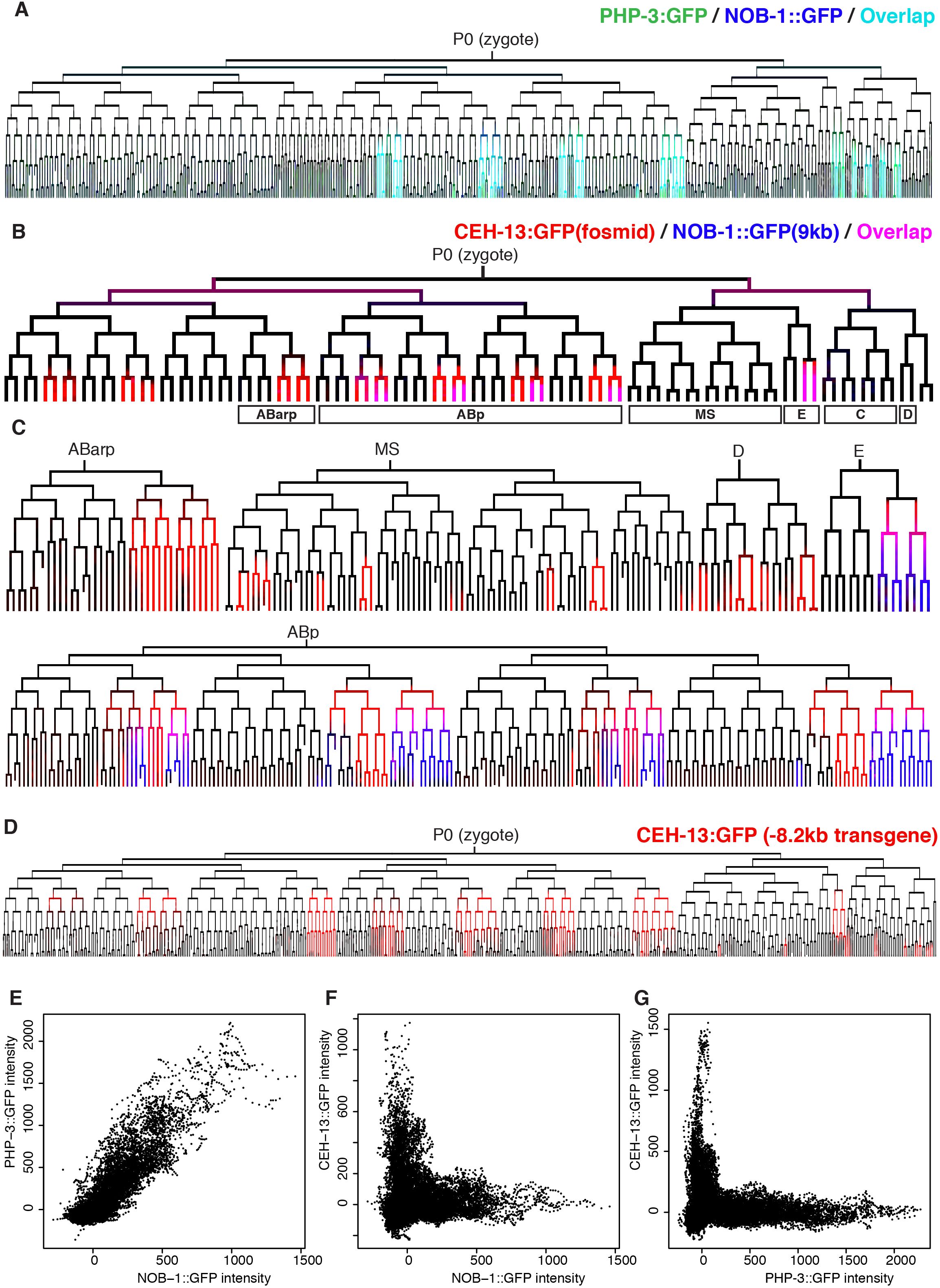
Hox gene are expressed in overlapping patterns. A) Overlap in the expression of NOB-1::GFP and PHP-3::GFP CRISPR knock-in alleles. Expression is nearly identical except *php-3* is expressed more consistently in ABplpapppp. B, C) Early (B) and late (C) overlap of a CEH-13::GFP fosmid translational reporter transgene with a -9kb NOB-1::GFP rescuing translational reporter transgene. The expression patterns are nearly identical to the CRISPR knock-in alleles except the 9kb NOB-1::GFP reporter has reduced expression in the C lineage. D) Expression pattern of a CEH-13::GFP transgene that contains 8.2 kb of upstream sequence plus the first intron. It lacks expression in the MS lineage compared to the other CEH- 13 reporters. E) Correlation between average CEH-13::GFP CRISPR and NOB-1::GFP CRISPR reporters for each cell during embryonic development. F) Correlation between average CEH- 13::GFP CRISPR and NOB-1::GFP CRISPR reporters for each cell during embryonic development. Note that no cells express high levels of both proteins. G) Correlation between average CEH-13::GFP and PHP-3::GFP intensity levels for each cell at each (1.5 minute) time point. Note that no cells express high levels of both proteins.

**Supplemental Figure 2:**
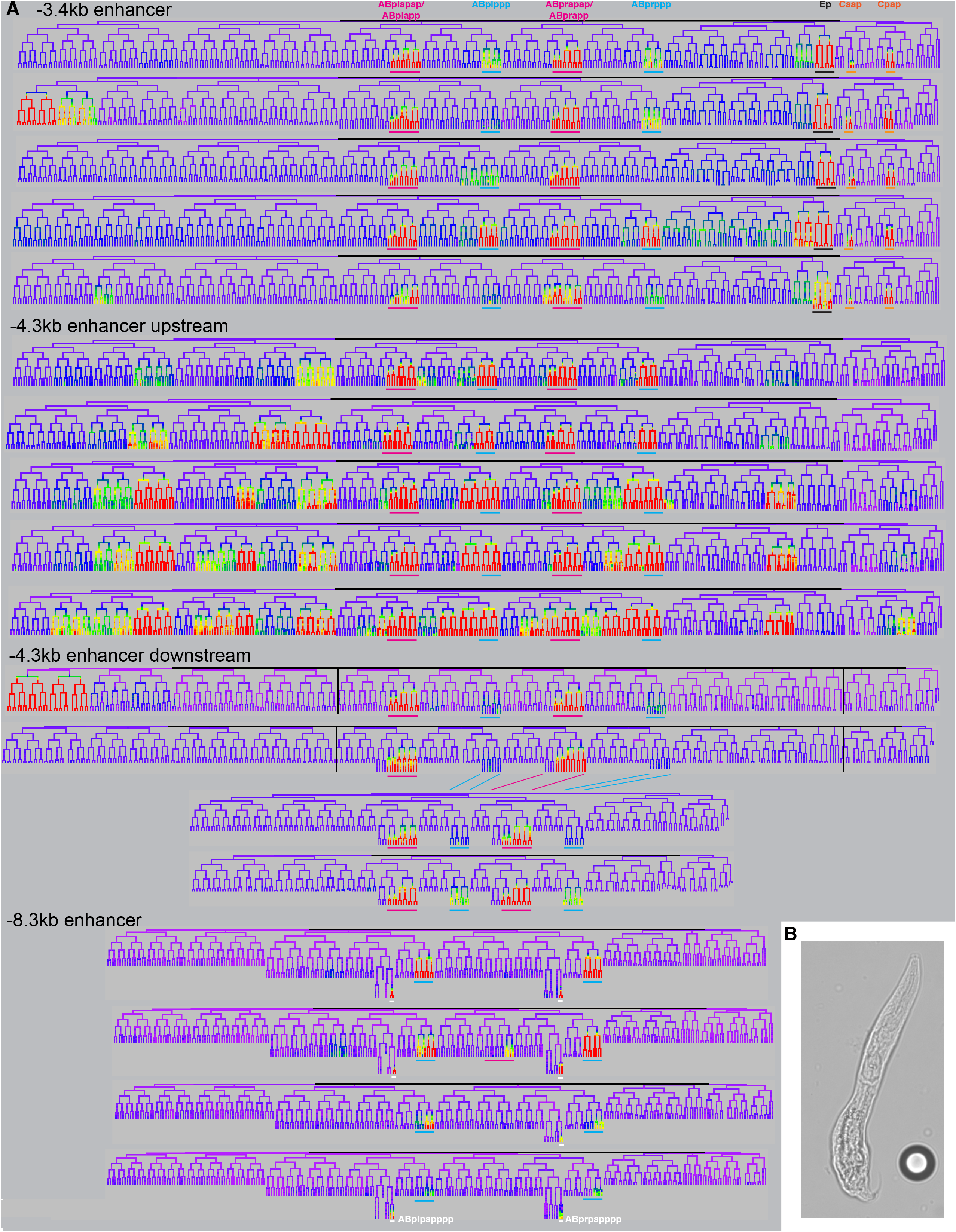
Some *nob-1* enhancer constructs drive variable ectopic expression. A) Representative examples showing the expression variability for each of the enhancers tested. Colored lines indicate the lineages of interest. Some lineages with no expression are shown as partial trees. B) Brightfield image of a hatched L1 larva carrying the -4.3kb enhancer reporter and showing the “no backend” phenotype characteristic of *nob-1/php- 3* mutants. Nearby bead is 20μm in diameter.

**Supplemental Figure 3:**
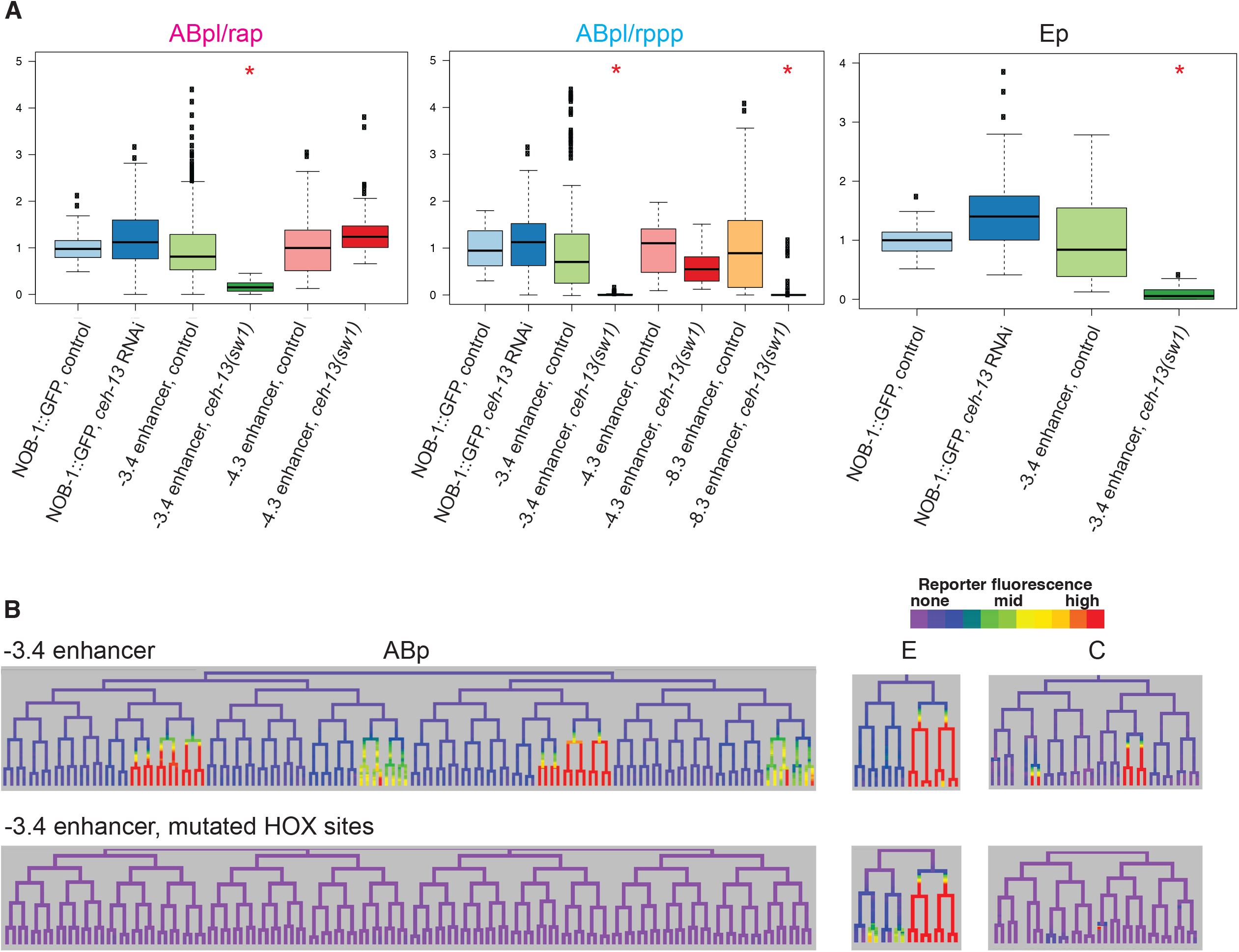
*ceh-13* regulates enhancers of *nob-1*. A) Graphs showing the effect of *ceh-13* RNAi on NOB-1::GFP rescuing transgene as well as the control values for the *nob-1* enhancer reporters (*sw1* mutant values same as reported in Fig. 3). Red * indicates p<0.05 in Wilcoxon Ranked sum test. B) Lineage trees showing expression of the wild-type -3.4kb enhancer and a version with mutated HOX sites.

**Supplemental Figure 4:**
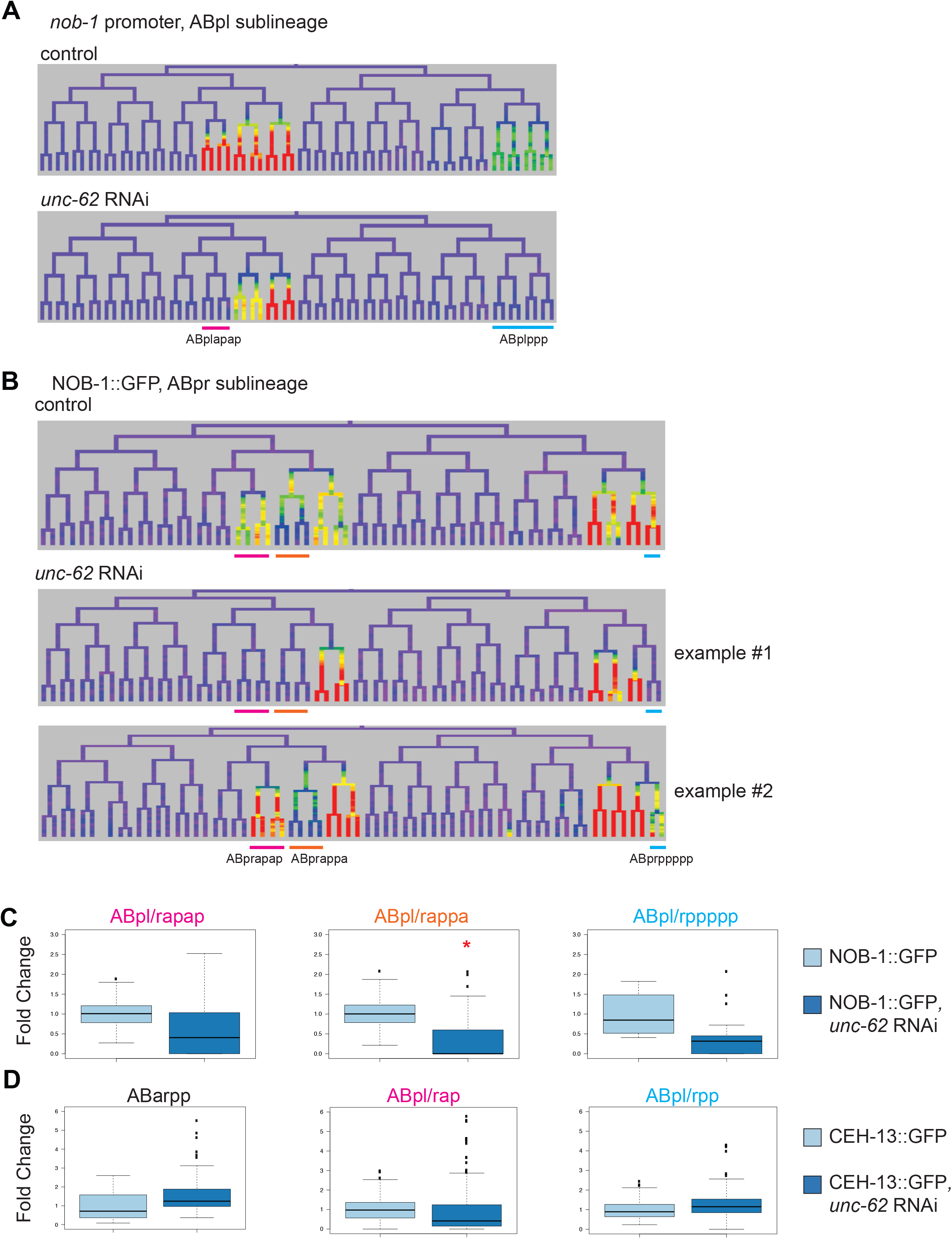
Hox co-factors affect *nob-1* expression in specific lineages. A) Tree showing effects of *unc-62i* on *nob-1* promoter transcriptional reporter expression in the ABpl lineage. Lineages with loss of expression are underlined. The ABp(l/r)apap lineage was most affected in 13/14 lineages tested. B) Trees showing the effect of *unc-62i* on NOB-1::GFP translational reporter expression, with two examples shown. Lineages affected are underlined. Graphs for all embryos tested (eight) for the same lineages shown in (B). Red * indicates p<0.05 in Wilcoxon Ranked sum test. D) Effect of *unc-62* RNAi on CEH-13::GFP fosmid reporter in lineages where the two genes are both expressed. No significant changes were detected.

**Supplemental Figure 5:**
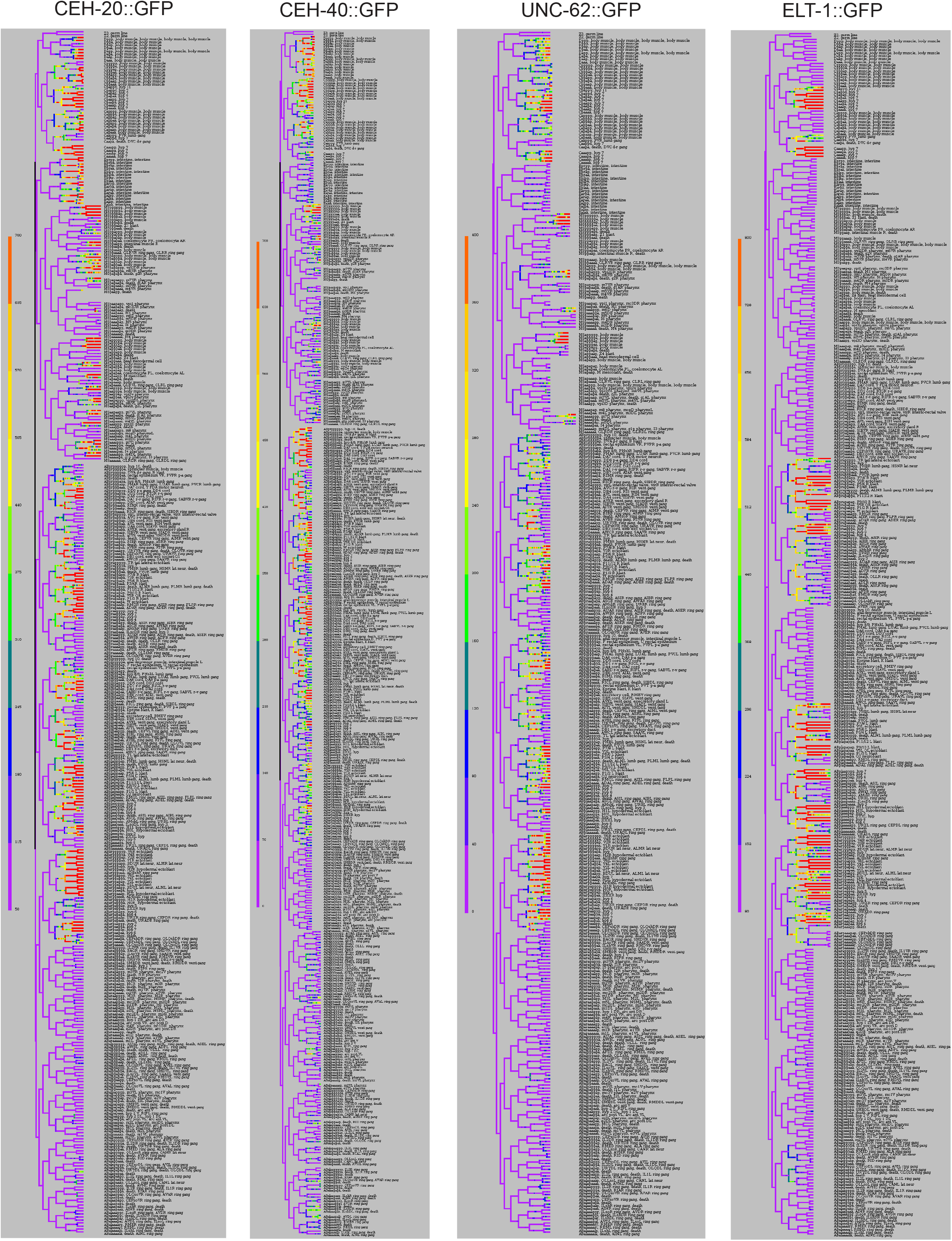
Full Lineages for *nob-1* regulators. Full lineages for CEH-20::GFP, CEH-40::GFP, UNC-62::GFP, and ELT-1::GFP, all fosmid translational reporters. All lineages are shown to at least the 350 cell stage - selected lineages are shown later to identify additional expression or dynamics.

**Supplemental Figure 6:**
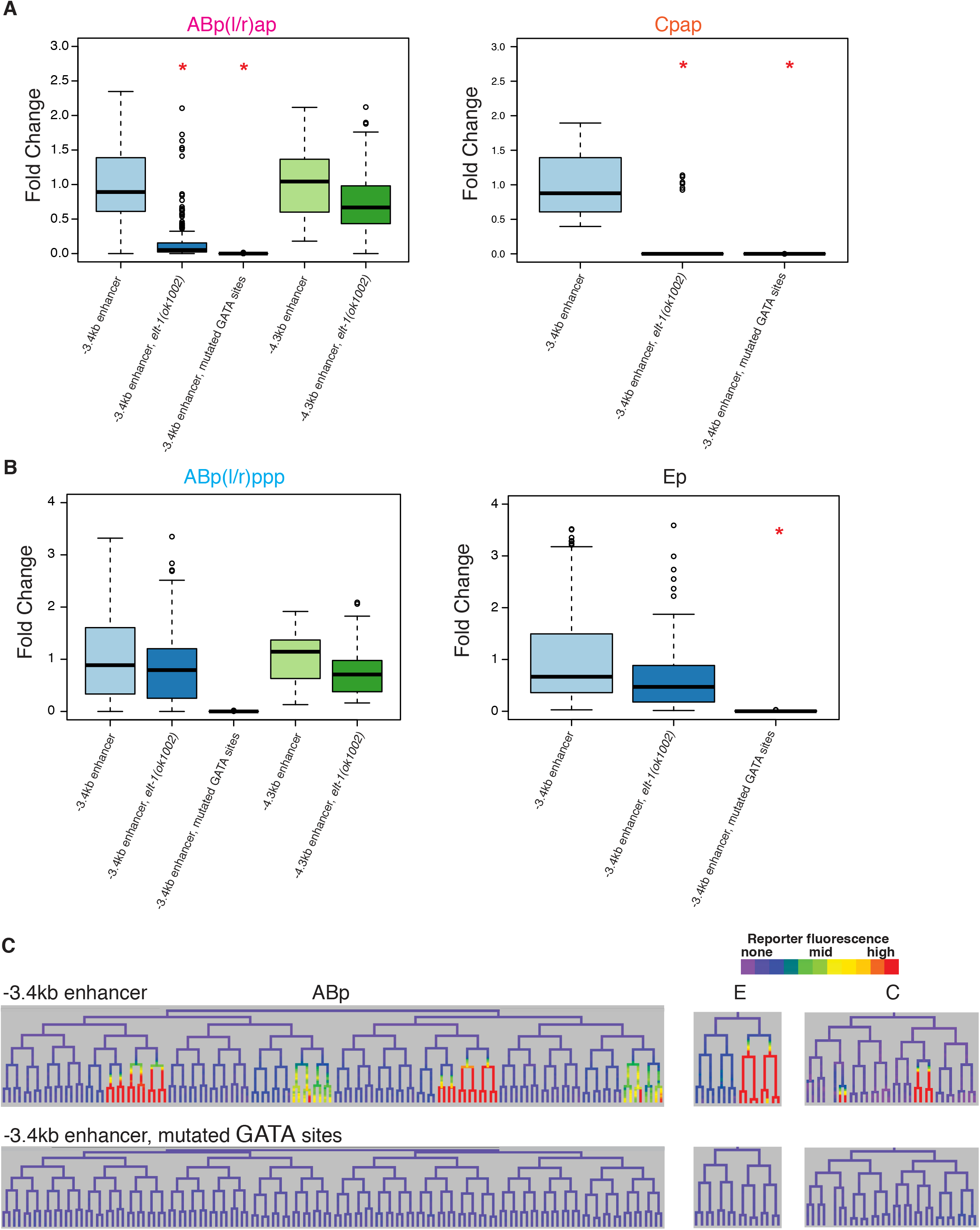
*nob-1* enhancer expression requires *elt-1*. A) Fold change values of *nob-1* enhancers in control and *elt-1* mutant conditions in the ABp(l/r)ap and Cpap lineages (if expressed in control). Mutant values are the same as reported in Fig. 5. Red * indicates p<0.05 in Wilcoxon Ranked sum test. B) Fold change values of *nob-1* enhancers in control and *elt-1* mutant conditions in the ABp(l/r)ppp and Ep lineages (if expressed in control) where ELT- 1::GFP is not expressed. Red * indicates p<0.05 in Wilcoxon Ranked sum test. C) Expression of the -3.4 kb enhancer and a version from which all GATA sites have been mutagenized in the ABp, E and C lineages. Note: the mutagenesis also disrupted one *ceh-20/40* predicted site, two *nob-1* predicted sites, and one *pop-1* predicted site, as these were fully overlapping with GATA sites.

**Supplemental Figure 7:**
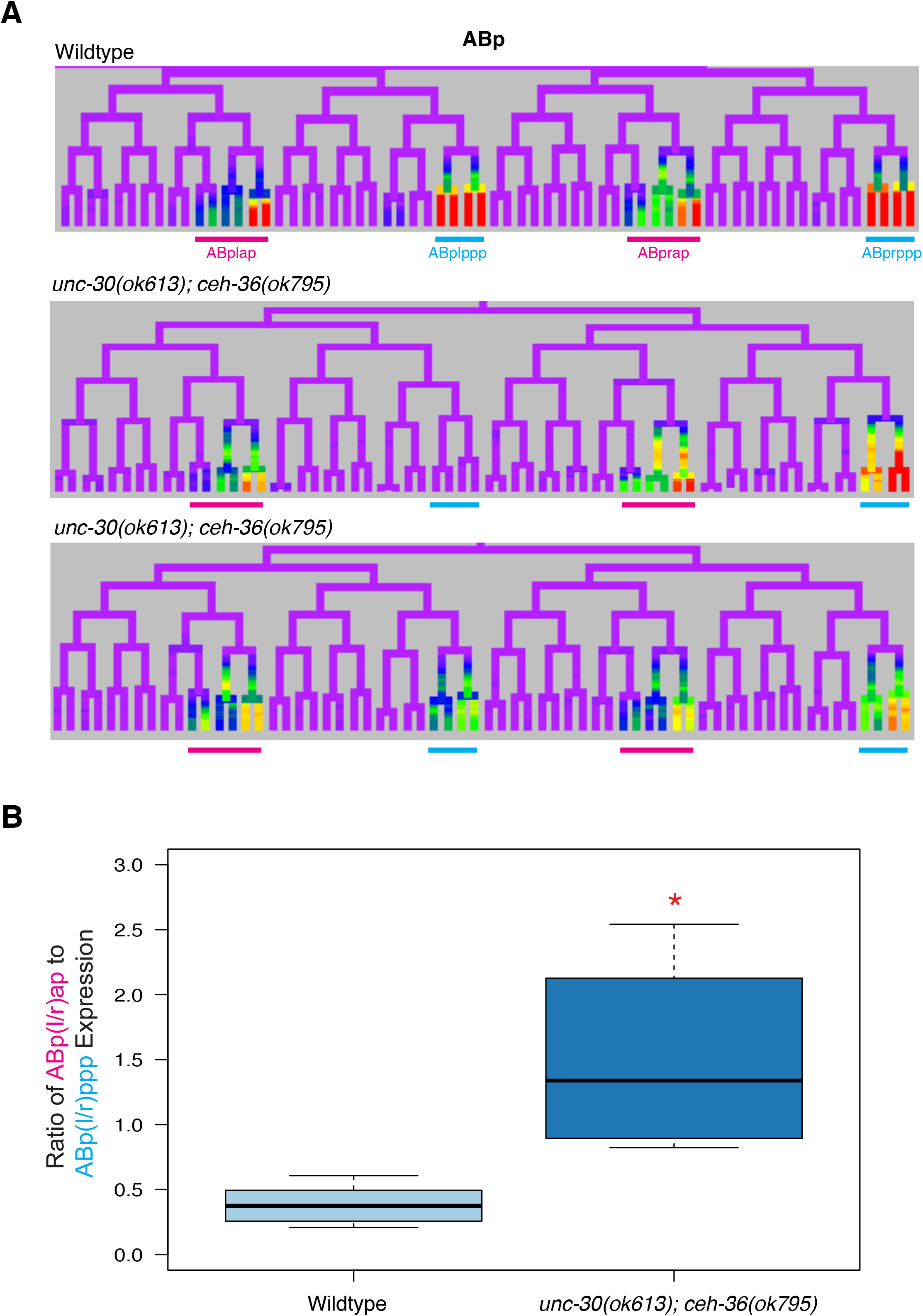
*ceh-36* and *unc-30* contribute to *nob-1* expression. A) Wild-type and *ceh-36(ok795);unc-30(ok613)* mutant embryos expressing the NOB-1::GFP reporter transgene. Expression is shown in the ABp lineage at the ∼200 cell stage. B) Boxplot showing the ratio of expression in the ABp(l/r)ap lineage to the ABp(l/r)ppp lineage for wild-type and *ceh- 36(ok795);unc-30(ok613)* mutant at the 350 cell stage for at least 4 embryos. Red * indicates p<0.05 in Wilcoxon Ranked sum test.

**Supplemental Figure 8:**
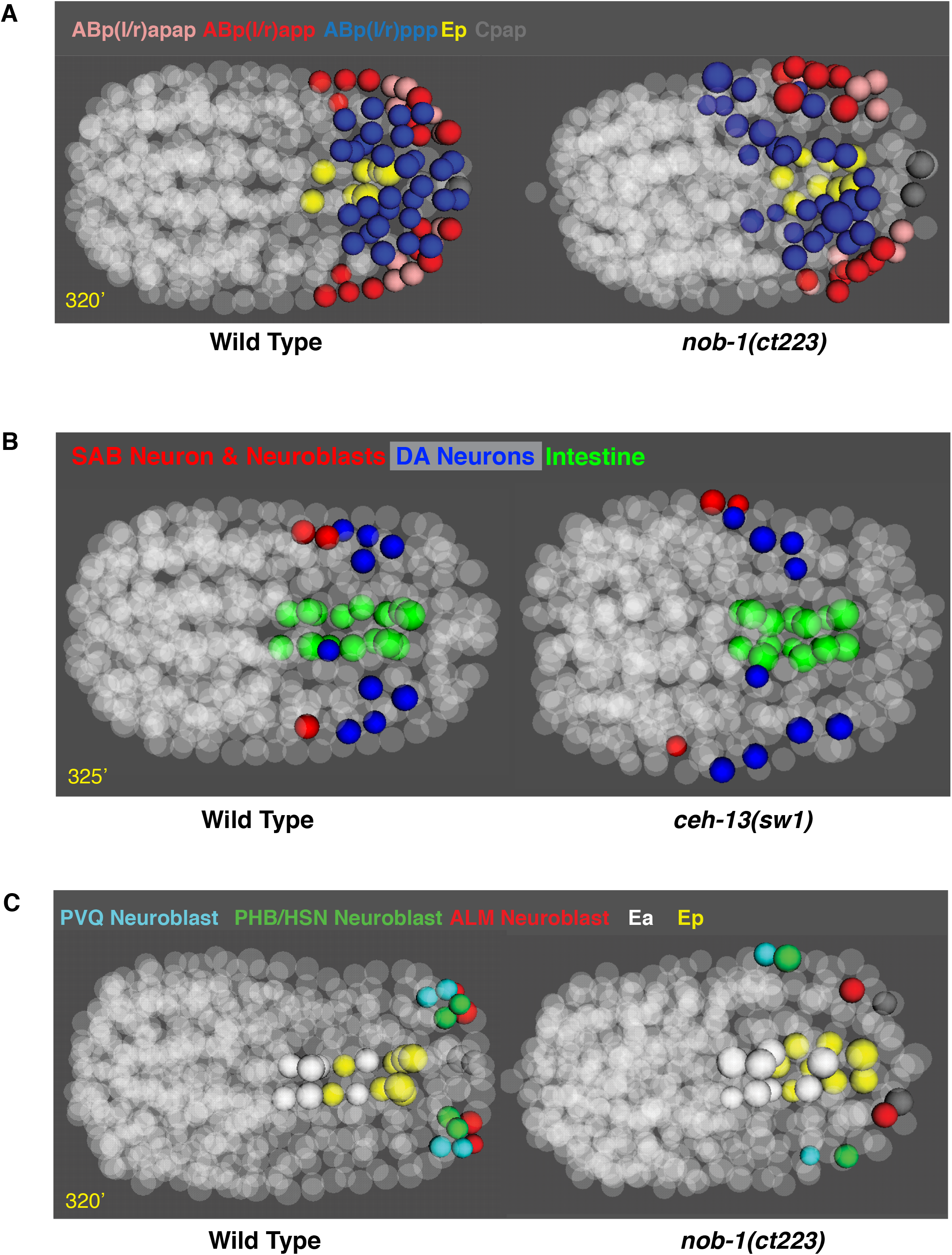
Defective positions of specific cells in *nob-1* mutants. A) The position of cells that normally express *nob-1* are highlighted in the context of the whole embryo at ∼320 minutes (∼570 cells, early morphogenesis). As compared to the wild-type average (left), the *nob-1(ct223)* mutant (right) cells are disorganized and displaced anteriorly, particularly ABp(l/r)ppp and some ABp(l/r)app cells (ABp(l/r)appp). B) The positions of the SAB neuron and neuroblasts (red) and DA motorneurons (blue) are shown relative to the intestine (green) in wild- type (left) and *ceh-13(sw1)* mutant (right), showing the anterior displacement of these cells. C) The positions of the neuroblasts that fail to divide the *nob-1(ct223)* mutant as compared to control (left), showing that these are dramatically mispositioned.

**Supplemental Figure 9:**
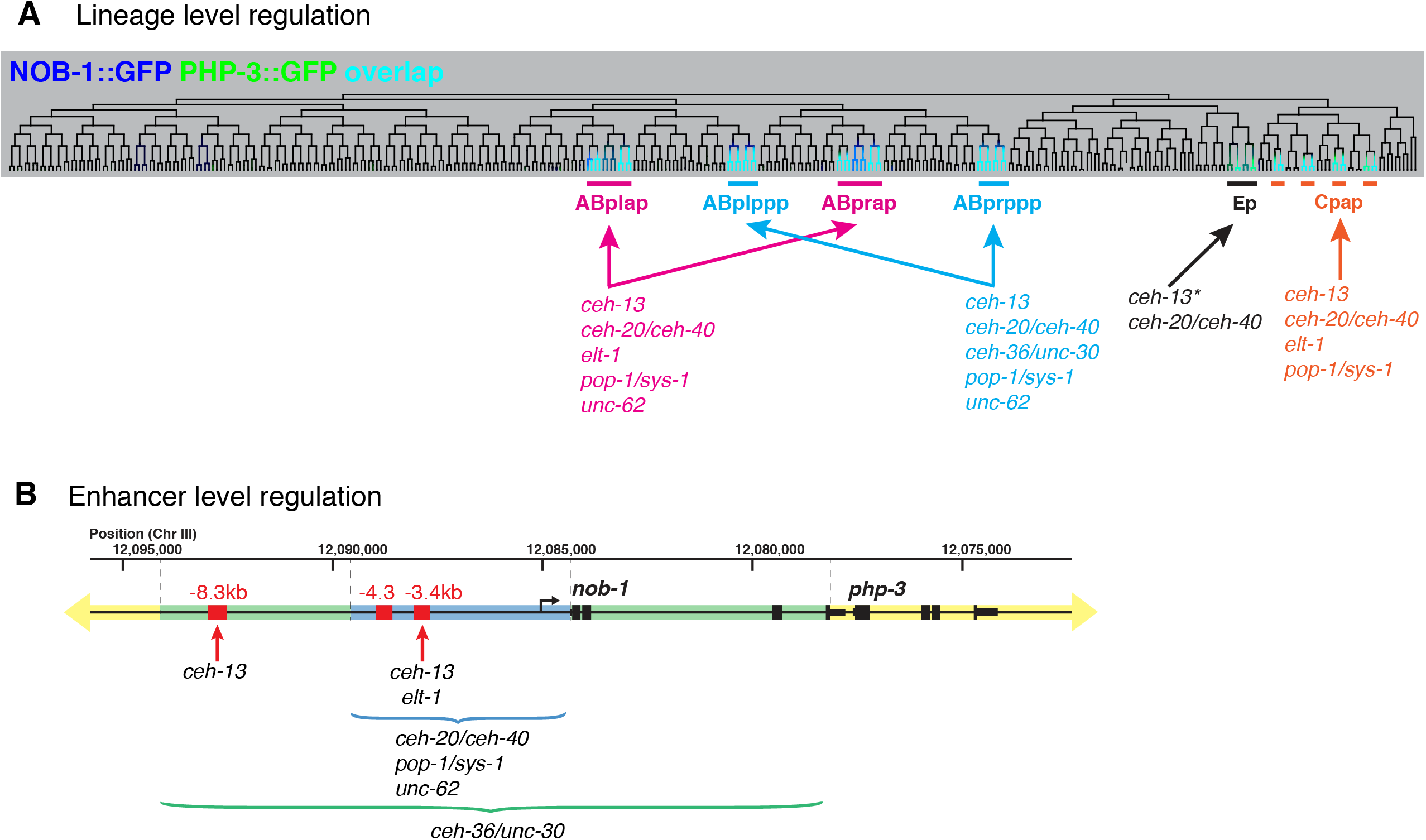
Regulation of *nob-1/php-3* at the lineage and enhancer levels. A) Diagram summarizing the transcriptional regulation of *nob-1/php-3* expression in the specific lineages noted. Asterisk indicates indirect regulation. Note: although *nob-1/php-3* are expressed in other lineages in C, only Cpap was possible to analyze with the transgenic reporters examined. Regulation by *pop-1/sys-1* from Zacharias et al., 2015 (Zacharias et al., 2015) B) Diagram summarizing enhancer-level transcriptional regulation of *nob-1/php-3* by the indicated factors. Specific enhancers regulated by *ceh-20/ceh-40*, *ceh-36/unc-30*, *pop-1/sys-1* and *unc-62* were not defined, but their activity can be localized to the regions marked by brackets. Based on our results, additional *cis*-regulatory elements likely exist within the blue, green and yellow regions (yellow encompasses the rest of the genome).

## Supplemental Tables found in SupplementalTables1-10.xlsx

Supplemental Table 1: Strain list

Supplemental Table 2: Tested enhancer sequences

Supplemental Tables 3-5: Motifs found in nob-1 enhancers -3.4kb, -4.3kb, -8.3kb

Supplemental Table 6: Cell averaged reporter values for all embryos analyzed

Supplemental Table 7: Number of embryos analyzed for each genotype/condition

Supplemental Tables 8-10: Cell division times for wild-type, *ceh-13(sw1),* and *nob-1(ct223)* embryos

## Supplemental Tables 11-13

Supplemental Table 11: Cell position values for control embryos

Supplemental Table 12: Cell position values for *ceh-13(sw1)* embryos

Supplemental Table 13: Cell position values for *nob-1(ct223)* embryos

## References Cited

1. Duboule D, Morata G. Colinearity and functional hierarchy among genes of the homeotic complexes. Trends Genet TIG. 1994 Oct;10(10):358–64.

2. Noordermeer D, Duboule D. Chromatin architectures and Hox gene collinearity. Curr Top Dev Biol. 2013;104:113–48.

3. Gaunt SJ. Hox cluster genes and collinearities throughout the tree of animal life. Int J Dev Biol. 2018;62(11–12):673–83.

4. Bürglin TR, Ruvkun G, Coulson A, Hawkins NC, McGhee JD, Schaller D, et al. Nematode homeobox cluster. Nature. 1991 Jun 27;351(6329):703.

5. Van Auken K, Weaver DC, Edgar LG, Wood WB. Caenorhabditis elegans embryonic axial patterning requires two recently discovered posterior-group Hox genes. Proc Natl Acad Sci U S A. 2000 Apr 25;97(9):4499–503.

6. Aboobaker A, Blaxter M. Hox gene evolution in nematodes: novelty conserved. Curr Opin Genet Dev. 2003 Dec;13(6):593–8.

7. Schaller D, Wittmann C, Spicher A, Müller F, Tobler H. Cloning and analysis of three new homeobox genes from the nematode Caenorhabditis elegans. Nucleic Acids Res. 1990 Apr 25;18(8):2033–6.

8. Bürglin TR, Ruvkun G. The Caenorhabditis elegans homeobox gene cluster. Curr Opin Genet Dev. 1993 Aug;3(4):615–20.

9. Weirauch MT, Yang A, Albu M, Cote AG, Montenegro-Montero A, Drewe P, et al. Determination and inference of eukaryotic transcription factor sequence specificity. Cell. 2014 Sep 11;158(6):1431–43.

10. Ferreira HB, Zhang Y, Zhao C, Emmons SW. Patterning of Caenorhabditis elegans posterior structures by the Abdominal-B homolog, egl-5. Dev Biol. 1999 Mar 1;207(1):215–28.

11. Harris J, Honigberg L, Robinson N, Kenyon C. Neuronal cell migration in C. elegans: regulation of Hox gene expression and cell position. Dev Camb Engl. 1996 Oct;122(10):3117–31.

12. Josephson MP, Chai Y, Ou G, Lundquist EA. EGL-20/Wnt and MAB-5/Hox Act Sequentially to Inhibit Anterior Migration of Neuroblasts in C. elegans. PloS One. 2016;11(2):e0148658.

13. Maloof JN, Whangbo J, Harris JM, Jongeward GD, Kenyon C. A Wnt signaling pathway controls hox gene expression and neuroblast migration in C. elegans. Dev Camb Engl. 1999 Jan;126(1):37–49.

14. Shemer G, Podbilewicz B. LIN-39/Hox triggers cell division and represses EFF-1/fusogen- dependent vulval cell fusion. Genes Dev. 2002 Dec 15;16(24):3136–41.

15. Takács-Vellai K, Vellai T, Chen EB, Zhang Y, Guerry F, Stern MJ, et al. Transcriptional control of Notch signaling by a HOX and a PBX/EXD protein during vulval development in C. elegans. Dev Biol. 2007 Feb 15;302(2):661–9.

16. Tihanyi B, Vellai T, Regos A, Ari E, Müller F, Takács-Vellai K. The C. elegans Hox gene ceh-13 regulates cell migration and fusion in a non-colinear way. Implications for the early evolution of Hox clusters. BMC Dev Biol. 2010 Jul 28;10:78.

17. Wang BB, Müller-Immergluck MM, Austin J, Robinson NT, Chisholm A, Kenyon C. A homeotic gene cluster patterns the anteroposterior body axis of C. elegans. Cell. 1993 Jul 16;74(1):29–42.

18. Yu H, Seah A, Sternberg PW. Re-programming of C. elegans male epidermal precursor fates by Wnt, Hox, and LIN-12/Notch activities. Dev Biol. 2010 Sep 1;345(1):1–11.

19. Zheng C, Jin FQ, Chalfie M. Hox Proteins Act as Transcriptional Guarantors to Ensure Terminal Differentiation. Cell Rep. 2015 Nov 17;13(7):1343–52.

20. Wittmann C, Bossinger O, Goldstein B, Fleischmann M, Kohler R, Brunschwig K, et al. The expression of the C. elegans labial-like Hox gene ceh-13 during early embryogenesis relies on cell fate and on anteroposterior cell polarity. Dev Camb Engl. 1997 Nov;124(21):4193–200.

21. Brunschwig K, Wittmann C, Schnabel R, Bürglin TR, Tobler H, Müller F. Anterior organization of the Caenorhabditis elegans embryo by the labial-like Hox gene ceh-13. Dev Camb Engl. 1999 Apr;126(7):1537–46.

22. Zhao Z, Boyle TJ, Liu Z, Murray JI, Wood WB, Waterston RH. A negative regulatory loop between microRNA and Hox gene controls posterior identities in Caenorhabditis elegans. PLoS Genet. 2010 Sep 2;6(9):e1001089.

23. Zacharias AL, Walton T, Preston E, Murray JI. Quantitative Differences in Nuclear β-catenin and TCF Pattern Embryonic Cells in C. elegans. PLoS Genet. 2015 Oct;11(10):e1005585.

24. Streit A, Kohler R, Marty T, Belfiore M, Takacs-Vellai K, Vigano M-A, et al. Conserved regulation of the caenorhabditis elegans labial/Hox1 gene ceh-13. Dev Biol. 2002;242(2):96–108.

25. Murray JI, Bao Z, Boyle TJ, Waterston RH. The lineaging of fluorescently-labeled Caenorhabditis elegans embryos with StarryNite and AceTree. Nat Protoc. 2006 Nov;1(3):1468–76.

26. Bao Z, Murray JI, Boyle TJ, Ooi S-L, Sandel MJ, Waterston RH. Automated cell lineage tracing in Caenorhabditis elegans. Proc Natl Acad Sci. 2006 Feb;103(8):2707–12.

27. Richards JL, Zacharias AL, Walton T, Burdick JT, Murray JI. A quantitative model of normal Caenorhabditis elegans embryogenesis and its disruption after stress. Dev Biol. 2013 Feb 1;374(1):12–23.

28. Santella A, Du Z, Bao Z. A semi-local neighborhood-based framework for probabilistic cell lineage tracing. BMC Bioinformatics. 2014 Jun 25;15:217.

29. Packer JS, Zhu Q, Huynh C, Sivaramakrishnan P, Preston E, Dueck H, et al. A lineage- resolved molecular atlas of C. elegans embryogenesis at single-cell resolution. Science. 2019 20;365(6459).

30. Kuntz SG, Schwarz EM, DeModena JA, De Buysscher T, Trout D, Shizuya H, et al. Multigenome DNA sequence conservation identifies Hox cis-regulatory elements. Genome Res. 2008 Dec;18(12):1955–68.

31. Ho MCW, Quintero-Cadena P, Sternberg PW. Genome-wide discovery of active regulatory elements and transcription factor footprints in Caenorhabditis elegans using DNase-seq. Genome Res. 2017 Dec;27(12):2108–19.

32. Sulston J. The embryonic cell lineage of the nematode Caenorhabditis elegans. Dev Biol. 1983 Nov;100(1):64–119.

33. Teng Y, Girard L, Ferreira HB, Sternberg PW, Emmons SW. Dissection of cis-regulatory elements in the C. elegans Hox gene egl-5 promoter. Dev Biol. 2004 Dec 15;276(2):476–92.

34. Coghlan A, Fiedler TJ, McKay SJ, Flicek P, Harris TW, Blasiar D, et al. nGASP--the nematode genome annotation assessment project. BMC Bioinformatics. 2008 Dec 19;9:549.

35. Gupta BP, Sternberg PW. The draft genome sequence of the nematode Caenorhabditis briggsae, a companion to C. elegans. Genome Biol. 2003;4(12):238.

36. Hillier LW, Miller RD, Baird SE, Chinwalla A, Fulton LA, Koboldt DC, et al. Comparison of C. elegans and C. briggsae Genome Sequences Reveals Extensive Conservation of Chromosome Organization and Synteny. PLOS Biol. 2007 Jul 3;5(7):e167.

37. Sternberg PW, Waterston RH, Spieth J, Eddy S, Wilson RK. Genome Sequence of Additional Caenorhabditis species: Enhancing the Utility of C. elegans as a Model Organism. Natl Hum Genome Res Inst White Pap. 2003 Oct 10;13.

38. Daugherty AC, Yeo RW, Buenrostro JD, Greenleaf WJ, Kundaje A, Brunet A. Chromatin accessibility dynamics reveal novel functional enhancers in C. elegans. Genome Res. 2017 Dec;27(12):2096–107.

39. Jänes J, Dong Y, Schoof M, Serizay J, Appert A, Cerrato C, et al. Chromatin accessibility dynamics across C. elegans development and ageing. eLife. 2018 Oct 26;7.

40. Murray JI, Boyle TJ, Preston E, Vafeados D, Mericle B, Weisdepp P, et al. Multidimensional regulation of gene expression in the C. elegans embryo. Genome Res. 2012 Jul;22(7):1282–94.

41. Niu W, Lu ZJ, Zhong M, Sarov M, Murray JI, Brdlik CM, et al. Diverse transcription factor binding features revealed by genome-wide ChIP-seq in C. elegans. Genome Res. 2011 Feb 1;21(2):245–54.

42. Araya CL, Kawli T, Kundaje A, Jiang L, Wu B, Vafeados D, et al. Regulatory analysis of the C. elegans genome with spatiotemporal resolution. Nature. 2014 Aug 28;512(7515):400–5.

43. Slattery M, Riley T, Liu P, Abe N, Gomez-Alcala P, Dror I, et al. Cofactor binding evokes latent differences in DNA binding specificity between Hox proteins. Cell. 2011 Dec 9;147(6):1270–82.

44. Abe N, Dror I, Yang L, Slattery M, Zhou T, Bussemaker HJ, et al. Deconvolving the recognition of DNA shape from sequence. Cell. 2015 Apr 9;161(2):307–18.

45. Van Auken K, Weaver D, Robertson B, Sundaram M, Saldi T, Edgar L, et al. Roles of the Homothorax/Meis/Prep homolog UNC-62 and the Exd/Pbx homologs CEH-20 and CEH-40 in C. elegans embryogenesis. Dev Camb Engl. 2002 Nov;129(22):5255–68.

46. Baugh LR, Wen JC, Hill AA, Slonim DK, Brown EL, Hunter CP. Synthetic lethal analysis of Caenorhabditis elegans posterior embryonic patterning genes identifies conserved genetic interactions. Genome Biol. 2005;6(5):R45.

47. Mace DL, Weisdepp P, Gevirtzman L, Boyle T, Waterston RH. A high-fidelity cell lineage tracing method for obtaining systematic spatiotemporal gene expression patterns in Caenorhabditis elegans. G3 Bethesda Md. 2013 May 20;3(5):851–63.

48. Narasimhan K, Lambert SA, Yang AWH, Riddell J, Mnaimneh S, Zheng H, et al. Mapping and analysis of Caenorhabditis elegans transcription factor sequence specificities. eLife. 2015 Apr 23;4.

49. Tintori SC, Osborne Nishimura E, Golden P, Lieb JD, Goldstein B. A Transcriptional Lineage of the Early C. elegans Embryo. Dev Cell. 2016 22;38(4):430–44.

50. Walton T, Preston E, Nair G, Zacharias AL, Raj A, Murray JI. The Bicoid class homeodomain factors ceh-36/OTX and unc-30/PITX cooperate in C. elegans embryonic progenitor cells to regulate robust development. PLoS Genet. 2015 Mar;11(3):e1005003.

51. Boeck ME, Boyle T, Bao Z, Murray J, Mericle B, Waterston R. Specific roles for the GATA transcription factors end-1 and end-3 during C. elegans E-lineage development. Dev Biol. 2011 Oct 15;358(2):345–55.

52. Jin Y, Hoskins R, Horvitz HR. Control of type-D GABAergic neuron differentiation by C. elegans UNC-30 homeodomain protein. Nature. 1994 Dec 22;372(6508):780–3.

53. Chang S, Johnston RJ, Hobert O. A transcriptional regulatory cascade that controls left/right asymmetry in chemosensory neurons of C. elegans. Genes Dev. 2003 Sep 1;17(17):2123–37.

54. Lanjuin A, VanHoven MK, Bargmann CI, Thompson JK, Sengupta P. Otx/otd homeobox genes specify distinct sensory neuron identities in C. elegans. Dev Cell. 2003 Oct;5(4):621– 33.

55. An JH, Vranas K, Lucke M, Inoue H, Hisamoto N, Matsumoto K, et al. Regulation of the Caenorhabditis elegans oxidative stress defense protein SKN-1 by glycogen synthase kinase-3. Proc Natl Acad Sci U S A. 2005 Nov 8;102(45):16275–80.

56. Bowerman B, Eaton BA, Priess JR. skn-1, a maternally expressed gene required to specify the fate of ventral blastomeres in the early C. elegans embryo. Cell. 1992 Mar 20;68(6):1061–75.

57. Gilleard JS, McGhee JD. Activation of hypodermal differentiation in the Caenorhabditis elegans embryo by GATA transcription factors ELT-1 and ELT-3. Mol Cell Biol. 2001 Apr;21(7):2533–44.

58. Ma X, Zhao Z, Xiao L, Xu W, Wang Y, Zhang Y, et al. Single-Cell Protein Atlas of Transcription Factors Reveals the Combinatorial Code for Spatiotemporal Patterning the C. elegans Embryo. bioRxiv. 2020 Jul 7;2020.06.30.178640.

59. Carroll SB. Evolution at two levels: on genes and form. PLoS Biol. 2005 Jul;3(7):e245.

60. Levine M. Transcriptional enhancers in animal development and evolution. Curr Biol CB. 2010 Sep 14;20(17):R754–763.

61. Cowing D, Kenyon C. Correct Hox gene expression established independently of position in Caenorhabditis elegans. Nature. 1996 Jul 25;382(6589):353–6.

62. Dupuy D, Li Q-R, Deplancke B, Boxem M, Hao T, Lamesch P, et al. A First Version of the Caenorhabditis elegans Promoterome. Genome Res. 2004 Oct 15;14(10b):2169–75.

63. Hunt-Newbury R, Viveiros R, Johnsen R, Mah A, Anastas D, Fang L, et al. High-throughput in vivo analysis of gene expression in Caenorhabditis elegans. PLoS Biol. 2007 Sep;5(9):e237.

64. Reece-Hoyes JS, Shingles J, Dupuy D, Grove CA, Walhout AJM, Vidal M, et al. Insight into transcription factor gene duplication from Caenorhabditis elegans Promoterome-driven expression patterns. BMC Genomics. 2007 Jan 23;8:27.

65. Charest J, Daniele T, Wang J, Bykov A, Mandlbauer A, Asparuhova M, et al. Combinatorial Action of Temporally Segregated Transcription Factors. Dev Cell. 2020 Nov 23;55(4):483–499.e7.

66. Hong J-W, Hendrix DA, Levine MS. Shadow enhancers as a source of evolutionary novelty. Science. 2008 Sep 5;321(5894):1314.

67. Frankel N, Davis GK, Vargas D, Wang S, Payre F, Stern DL. Phenotypic robustness conferred by apparently redundant transcriptional enhancers. Nature. 2010 Jul 22;466(7305):490–3.

68. Perry MW, Boettiger AN, Bothma JP, Levine M. Shadow enhancers foster robustness of Drosophila gastrulation. Curr Biol CB. 2010 Sep 14;20(17):1562–7.

69. Maduro MF. Gut development in C. elegans. Semin Cell Dev Biol. 2017 Jun;66:3–11.

70. Sarov M, Murray JI, Schanze K, Pozniakovski A, Niu W, Angermann K, et al. A genome-scale resource for in vivo tag-based protein function exploration in C. elegans. Cell. 2012 Aug 17;150(4):855–66.

71. Green JL, Inoue T, Sternberg PW. Opposing Wnt pathways orient cell polarity during organogenesis. Cell. 2008 Aug 22;134(4):646–56.

72. Fire A, Harrison SW, Dixon D. A modular set of lacZ fusion vectors for studying gene expression in Caenorhabditis elegans. Gene. 1990 Sep 14;93(2):189–98.

73. Wei X, Potter CJ, Luo L, Shen K. Controlling gene expression with the Q repressible binary expression system in Caenorhabditis elegans. Nat Methods. 2012 Mar 11;9(4):391–5.

74. Bao Z, Murray JI. Mounting Caenorhabditis elegans Embryos for Live Imaging of Embryogenesis. Cold Spring Harb Protoc. 2011 Sep 1;2011(9):pdb.prot065599.

75. Boyle TJ, Bao Z, Murray JI, Araya CL, Waterston RH. AceTree: a tool for visual analysis of Caenorhabditis elegans embryogenesis. BMC Bioinformatics. 2006 Jun 1;7:275.

76. Murray JI, Bao Z, Boyle TJ, Boeck ME, Mericle BL, Nicholas TJ, et al. Automated analysis of embryonic gene expression with cellular resolution in C. elegans. Nat Methods. 2008 Jun;5(8):703–9.

